# In situ pulse dispersion estimation via third-harmonic generation interferometric autocorrelation for multiphoton microscopy

**DOI:** 10.64898/2026.08.04.742801

**Authors:** Liam Shaughnessy, Lauren Vannell, Sulekh Fernando-Peiris, Cristina Rodríguez

**Author notes:** Correspondence (C.R.). These authors contributed equally to this work (Liam Shaughnessy, Lauren Vannell).

## Abstract

Three-photon microscopy enables deep-tissue imaging but is highly sensitive to excitation pulse quality, since three-photon excitation efficiency scales with the cube of instantaneous intensity. Group-delay dispersion (GDD) and third-order dispersion (TOD) accumulated through the laser and microscope optics can substantially reduce peak intensity at the focal plane, yet these quantities are rarely measured where imaging occurs. Here, we use third-harmonic generation (THG) interferometric autocorrelation, together with Dispersion Look-Up-Table Estimation (D-LUTE) and a joint two-measurement fitting procedure validated on synthetic data, to estimate baseline GDD and TOD directly at the objective focal plane, requiring only a compact autocorrelator module added to the microscope beam path. Applying this method at 1300 nm and 1600 nm excitation across two microscope systems equipped with different units of the same laser model, we find pulse durations 1.4-to 1.7-fold longer than the transform limit at every condition, driven predominantly by TOD, which varied by roughly 2.7-fold between the two systems. Using the endogenous THG signal from myelinated fibers, we further demonstrate in vivo pulse characterization in the mouse brain, finding no measurable broadening between the tissue surface and a depth of nearly a millimeter. This low-cost, easily implemented approach enables routine, in situ pulse monitoring across multiphoton microscopy platforms.

## 1 INTRODUCTION

Three-photon (3P) microscopy (3PM) has emerged as a powerful tool for deep-tissue biological imaging, extending the penetration depth achievable with two-photon (2P) microscopy (2PM) by taking advantage of longer excitation wavelengths in the near-infrared (NIR-II, ∼1300 and ∼1700 nm) windows, where tissue scattering and absorption are reduced^1,2^. This has enabled minimally invasive, high-resolution imaging of structures and dynamics previously inaccessible to conventional multiphoton microscopy, transforming investigation across a range of biological systems^3^, from deep cortical and subcortical brain regions to other historically inaccessible, highly scattering tissues^4^.

Because 3P excitation efficiency scales with the cube of instantaneous intensity, 3PM is markedly more sensitive to the fidelity of the excitation pulse than 2PM. For a pulse of fixed energy, this cubic dependence means efficiency scales as the inverse square of pulse duration when dispersion simply stretches the pulse in time without altering its shape; more generally, efficiency depends on the full temporal profile. Consequently, any temporal distortion accumulated as the pulse propagates through the laser, microscope optics, and objective – most notably group-delay dispersion (GDD) and third-order dispersion (TOD) – reduces peak intensity and, in turn, imaging depth and signal brightness. TOD is particularly consequential for the broad-bandwidth pulses used in 3PM, distorting the pulse asymmetrically in ways not addressed by dispersion-compensation schemes designed primarily for GDD^5^.

Several techniques exist for characterizing the temporal and spectral phase of ultrashort pulses, and each relies on some nonlinear-optical signal – typically second-harmonic generation (SHG) or sum-frequency mixing – to gate or interfere the pulse against a delayed replica of itself. Frequency-resolved optical gating (FROG)^6^, spectral phase interferometry for direct electric-field reconstruction (SPIDER)^7^, and dispersion-scan (d-scan)^8^ resolve a pulse’s amplitude and phase unambiguously by adding an additional measurement dimension, but at the cost of increased experimental complexity. Intensity and interferometric autocorrelation, by contrast, are simpler and more widely used for routine pulse monitoring, requiring only a straightforward optical setup that can be readily performed at a microscope’s focal plane. However, reconstructing a pulse’s full amplitude and phase from a single autocorrelation trace alone is fundamentally ambiguous: different pulses can produce nearly identical autocorrelation traces and power spectra^9^. Early approaches combined the pulse spectrum with interferometric autocorrelation measurements to iteratively reconstruct the full field^10,11^, but they suffered from convergence issues without good initial guesses. One strategy for resolving the residual ambiguity is to introduce a second measurement with a known dispersive element added to the beam path, for example to determine the time-direction (sign) of asymmetric pulses from paired autocorrelation measurements^12^. This approach was recently applied within 3PM to resolve the direction-of-time ambiguity inherent to SHG-FROG, determining the sign of both GDD and TOD via measurements at the laser output, prior to propagation through the microscope^5^.

A complementary strategy, advantageous for microscopy, uses the sample itself as the autocorrelation nonlinearity, performing the measurement in situ at the focal plane. Previous work used second-order autocorrelation via 2P fluorescence to obtain absolute 2P cross-sections without prior knowledge of the pulse shape^13^, later extended to measure the GDD of high-NA objectives via the same 2P signal^14^. A third-order interferometric autocorrelator using 3P fluorescence from exogenous fluorophores has similarly been used to estimate GDD in a 3P microscope^15^, and to determine third-order temporal coherence for extracting 3P cross-sections via joint second- and third-order fluorescence autocorrelation^16^.

Third-harmonic generation (THG) offers a convenient nonlinear signal for such autocorrelation measurements. THG is a third-order nonlinear optical process^17,18^ in which three incoming excitation photons are coherently converted into a single photon at three times the energy, corresponding to one-third the wavelength of the excitation. For a tightly focused beam in a homogeneous medium, the generated THG field vanishes due to the Gouy phase shift^19^. However, this symmetry is broken at a material interface, where a net THG signal is generated. The process works efficiently using ordinary, inexpensive materials rather than specialized nonlinear crystals, while providing improved sensitivity to the pulse shape from its cubic field dependence. This same interface selectivity is also what makes THG a widely used label-free contrast mechanism in biological imaging, generated efficiently at lipid-rich structures such as myelin sheaths, cell membranes, and other interfaces within tissue^20,21^.

As a coherent, energy-conserving parametric process that proceeds through virtual states, THG generates no real excited-state population, avoiding the photobleaching and saturation that limit fluorescence-based measurements over extended acquisition and can bias the trace if bleaching progresses over the scan. This advantage has also been exploited for pulse characterization: THG generated at the surface of an ordinary glass slide was used both as the nonlinearity for interferometric and intensity autocorrelation of ultrashort pulses^22^, and in a full FROG geometry, for retrieval of the pulse’s spectral phase without time-direction ambiguity^23^. This was later extended to the focus of a microscope objective, enabling collinear, background-free THG-FROG retrieval of the pulse’s amplitude and phase at the objective focal plane^24^. However, this proof-of-concept retrieval requires spectrally resolved measurements that add experimental complexity, limiting its adoption for routine use.

A simpler, in situ approach to recovering a pulse’s baseline dispersion directly at the focal plane remains desirable. Here, we take advantage of THG’s interface selectivity to integrate a third-order interferometric autocorrelator directly into a THG microscope. Built from inexpensive off-the-shelf components, the module fits readily into the beam path of any multiphoton microscope capable of detecting the resulting THG signal, requiring minimal alignment and enabling in situ pulse characterization at the objective focal plane. From the resulting THG interferometric autocorrelation (THG-IAC) traces, acquired with and without added dispersive elements, we recover the excitation pulse’s baseline GDD and TOD in two steps: Dispersion Look-Up-Table Estimation (D-LUTE), which uses the measured change between the two conditions to provide physically informed initial estimates, followed by a joint fit of the paired trace envelopes, applied at both the 1300 nm and 1600 nm excitation windows across two independent microscope systems. Finally, we show that this module can be used for pulse characterization directly within intact, scattering biological tissue in vivo, taking advantage of the endogenous THG signal generated by myelin in the mouse brain.

## 2 EXPERIMENTAL SETUP AND METHODS

### 2.1 Microscope description

Fig. 1 provides an overview of the custom-built THG microscope, including the optical layout of the excitation, scanning, and autocorrelation module (Fig. 1a,b), the excitation point-spread function (PSF) (Fig. 1c), the laser spectrum (Fig. 1d), and the power dependence of the THG signal confirming its third-order origin (Fig. 1e).

**Figure 1.**
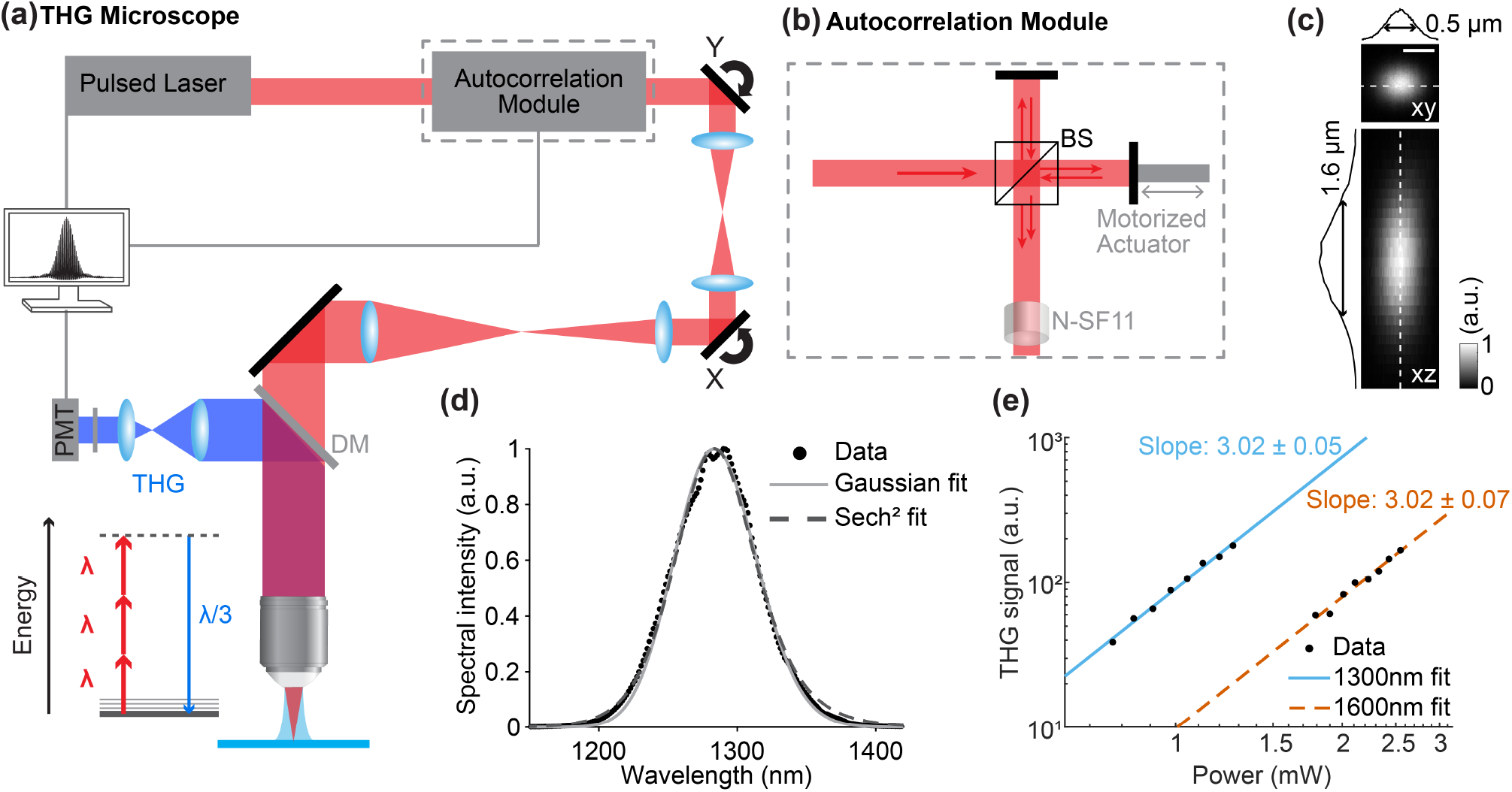
THG microscope and autocorrelation module. **(a)** Main components of the THG microscope and autocorrelation module; X, Y, galvanometers; DM, dichroic mirror; PMT, photomultiplier tube. For simplicity, only one of the two detection channels used in this study is shown. Bottom left: energy-level diagram of THG. **(b)** Detailed schematic of the autocorrelation module. BS, beamsplitter; N-SF11, glass rod. **(c)** Lateral (XY) and axial (XZ) THG images of a 200-nm-diameter gold bead at 1300 nm excitation, with measured FWHM (0.5 µm lateral, 1.6 µm axial). Post-objective power: 0.5 mW. Scale bar, 0.5 µm. **(d)** Measured laser spectrum at 1300 nm (dots), with Gaussian (solid) and sech^2^ (dashed) fits. **(e)** THG signal as a function of excitation power at 1300 nm and 1600 nm, generated at a water-glass interface at the focal plane; solid and dashed lines show power-law fits at 1300 nm and 1600 nm, respectively (slopes indicated), confirming the expected cubic dependence.

Excitation was provided by a tunable femtosecond laser (Cronus 3P, Light Conversion). The system incorporates an optical parametric amplifier (OPA) pumped by a 40-W, 1035 nm femtosecond laser operating at a 1 MHz repetition rate, producing a tunable output in the NIR-II range (1250-1800 nm). At 1300 nm, the primary wavelength used in this study, the laser delivered 1.2 W of average power (1.2 µJ per pulse). The laser spectrum was measured using a spectrometer (Sphere Photonics d-vision, Axiom Optics) with a spectral range of 950-1680 nm and a resolution of 2 nm. At 1300 nm, the measured center wavelength was 1284.4 nm, with a spectral width of approximately 69.6 nm (Fig. 1d). In comparison, at 1600 nm, the laser delivered 0.8 W of average power (0.8 µJ per pulse), with a measured center wavelength of 1583 nm and a spectral width of approximately 66.1 nm.

An internal two-prism compressor integrated into the laser system was used to minimize GDD and achieve the shortest achievable pulse duration at the focal plane of the microscope objective. After optimizing the compressor settings, pulse durations of 55 fs at 1300 nm and 77 fs at 1600 nm were measured at the focal plane of the microscope objective using our custom-built THG interferometric autocorrelator (Sections 3.2 and 3.3).

Two-dimensional raster scanning was performed using a pair of galvanometric mirrors optically conjugated to each other via a pair of scan lenses (Thorlabs, SL50-3P, f = 50 mm) and to the back pupil plane of a water-immersion objective (Olympus XLPLN25XWMP2, 25×, NA 1.05) through an additional scan lens and tube lens pair (Thorlabs, SL50-3P, f = 50 mm and TTL200MP, f = 200 mm). The measured lateral and axial resolutions were 0.5 µm and 1.6 µm, respectively (Fig. 1c). For axial translation of the laser focus, the objective was mounted on a piezoelectric stage (P-725.4CDE2 PIFOC; Physik Instrumente). The same objective epi-collected the THG signal, which was separated from the excitation light and directed into the detection module by a dichroic mirror (FF705-Di02-25×36, Semrock). The signal was further split into two detection channels using a dichroic mirror (FF496-SDi01-25×36, Semrock) and detected by GaAsP photomultiplier tubes (PMTs; H16201P-40-02, Hamamatsu) after going through bandpass filters to isolate the THG generated from 1300 nm (FF01-433/24-25, Semrock) and 1600 nm excitation (FF01-531/46-30-D, Semrock). Image acquisition was controlled using ScanImage (MBF Bioscience). Power-law fits to the THG signal generated at a water-glass interface as a function of excitation power confirmed the expected cubic dependence at both wavelengths (Fig. 1e).

N-SF11 glass rods (Fig. 1b) were used to introduce a known dispersion increment into the beam path for dispersion estimation (Section 2.6). Using the Sellmeier equation for N-SF11^25^ at the measured center wavelength of each dataset, the calculated GDD and TOD contributed by the rods were 1584 fs^2^ and 2879 fs^3^ for a single 20 mm rod at 1284.4 nm excitation wavelength, and 1815 fs^2^ and 13,581 fs^3^ for three 20 mm rods (60 mm total) at 1583 nm excitation wavelength.

### 2.2 Autocorrelation module

A home-built Michelson-type interferometer was integrated into the microscope beam path to enable direct measurement of laser pulse duration at the focal plane of the microscope objective (Fig. 1b; Supplementary Fig. S1, Supplementary Note S1). The excitation beam first passed through a 50:50 non-polarizing BK7 beamsplitter (CCM5-BS018, Thorlabs), which divided it into two arms. One arm contained a mirror mounted on a motorized actuator (Z812B, Thorlabs) driven by a DC servo controller (KDC101, Thorlabs), enabling precise control of the relative time delay between the two pulse replicas. After reflection, the two beams were recombined at the beamsplitter and directed back into the microscope optical path.

To maintain collinearity, each arm was carefully aligned by blocking one path at a time and verifying spatial overlap at multiple downstream locations using a beam profiler (BP209-IR2, Thorlabs). Optical power in the two arms was measured to differ by less than 1%, avoiding amplitude asymmetries that could otherwise distort the interferometric autocorrelation trace. Representative THG-IAC traces acquired with the module deliberately misaligned or with unequal arm powers are shown in Supplementary Fig. S2. The recombined beam was tightly focused onto a glass coverslip using a high-NA objective, and the resulting THG signal from the water-glass interface was epi-collected to generate THG interferometric autocorrelation (THG-IAC) traces.

The breadboard-mounted module was designed for straightforward insertion and removal from the beam path. A set of flip mirrors at the module entrance and exit allowed the interferometer to be bypassed entirely, enabling standard microscope operation without disturbing downstream alignment. Using the Sellmeier equation for BK7^26^, the beamsplitter is calculated to introduce a double-pass dispersion of GDD ≈ 139 fs^2^, TOD ≈ 3237 fs^3^ at 1284.4 nm and GDD ≈ −1155 fs^2^, TOD ≈ 6812 fs^3^ at 1583 nm. All baseline GDD and TOD values reported in Section 3 are those measured with the module in the beam path and therefore include this contribution; the dispersion present during standard imaging, with the module bypassed, is obtained by subtracting it.

### 2.3 Theoretical framework for THG-based interferometric autocorrelations

THG generated at a material interface provides a convenient signal source for third-order interferometric autocorrelation measurements. When two identical, collinear pulses, separated by a time delay τ, produced by our autocorrelator module (Fig. 1b), overlap at the interface, the resulting THG intensity scales with the sixth power of the combined electric field amplitude, |E|^6^. This strong nonlinear sensitivity to temporal overlap enables in situ characterization of femtosecond pulse duration at the microscope focal plane.

To model third-order interferometric autocorrelations, we represent the excitation pulses as electric fields of the form:

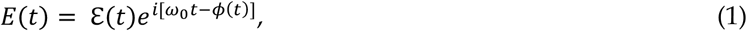

where *ω*_0_ is the carrier frequency, *ϕ*(*t*) is the time-dependent phase, and ε(*t*) is the real-valued pulse envelope. A common envelope model is the Gaussian pulse:

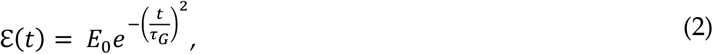

where 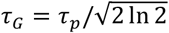 ln 2 and *τ*_*p*_ is the pulse duration, defined as the full width at half maximum (FWHM) of the intensity profile |ε(*t*)|^2^. Other shapes, such as the hyperbolic secant (sech), are used for sources with different spectral profiles; the appropriate envelope model depends on the specific laser source and should be verified against the measured spectrum.

While the time-domain formulation describes the pulse shape directly, dispersion effects are more naturally analyzed in the frequency domain, where the spectral phase determines the temporal profile. The frequency-domain field is obtained via the Fourier transform:

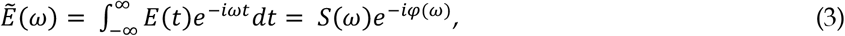

where *S*(*ω*) is the spectral amplitude and *φ*(*ω*) is the spectral phase. The spectral phase can be expanded as a Taylor series around the carrier frequency *ω*_0_:

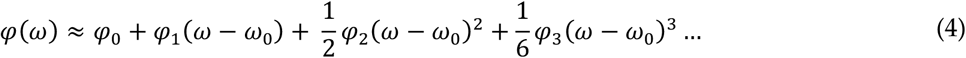

Here, *φ*_0_ represents a constant offset to the spectral phase (i.e., the carrier-envelope phase), *φ*_1_ corresponds to group delay, *φ*_2_ is the GDD, and *φ*_3_ is the TOD. The zeroth- and first-order terms affect only absolute phase and arrival time, leaving pulse width unchanged; higher-order terms distort the temporal profile – GDD produces symmetric broadening and linear chirp, while TOD introduces asymmetric pre- and post-pulses. These effects grow increasingly significant for broadband pulses, such as those used in this study.

To simulate dispersion, we apply this phase expansion to the frequency-domain field:

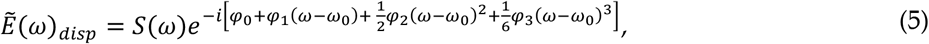

then recover the time-domain field by inverse Fourier transform, 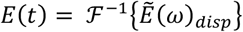. In what follows, *E*(*t*) denotes this dispersed field. Unlike a Gaussian pulse, a sech pulse or any more complex pulse shape has no closed-form solution once GDD is applied, and the problem becomes further complicated with TOD or higher-order terms. In general, the transformation must be carried out numerically using fast Fourier transforms (FFT).

Autocorrelation techniques are commonly used to characterize such pulses experimentally, measuring the nonlinear signal produced by the temporal overlap of two time-delayed pulse replicas. The nth-order intensity autocorrelation is defined as^27^:

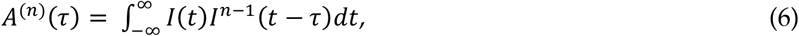

where *I*(*t*) is the pulse intensity, *τ* is the time delay, and *n* indicates the order of the nonlinear process involved. Intensity autocorrelation is simple to implement and is widely used for estimating pulse width; however, it provides no information about the spectral phase – including GDD and TOD.

Interferometric autocorrelations overcome this limitation by incorporating the full electric field: because the two pulse replicas add coherently before the nonlinear interaction, the resulting trace retains information about the pulse’s phase, including distortions introduced by GDD and TOD. The general expression for the nth-order interferometric autocorrelation is^27^:

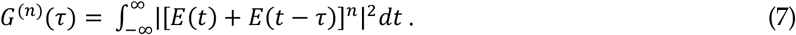

Using THG (*n* = 3) as the nonlinear mechanism, we use the third-order interferometric autocorrelation:

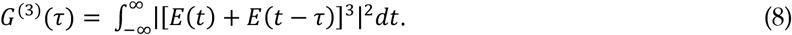

This sixth-order dependence on the electric field makes the THG signal highly sensitive to the degree of temporal overlap, enabling high-resolution pulse duration measurements. Expansion of Eq. (8) also produces interference terms that oscillate at the carrier frequency and its second and third harmonics, giving rise to the high-frequency fringes in the interferometric autocorrelation trace. Spectral filtering removes these fringes, leaving the intensity autocorrelation, *A*^(3)^(*τ*), on a constant background.

### 2.4 Simulated effects of dispersion on THG interferometric autocorrelation traces

To visualize how GDD and TOD affect THG-IAC traces, we numerically simulated the THG signal generated by interference of two time-delayed femtosecond pulse replicas.

To select an appropriate spectral model, we compared Gaussian and sech^2^ fits to the measured spectrum at 1300 nm, as well as their downstream effect on simulated autocorrelation-derived quantities (Supplementary Note S2). Although the two fits were of comparable quality, the Gaussian model more closely reproduced the quantities computed directly from the measured spectrum and was adopted for all simulations in this work.

Using the Gaussian fit to the measured spectrum (*λ*_0_ = 1284.4 nm, Δλ = 69.6 nm), we obtain a transform-limited pulse duration of 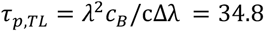fs, with *c*_*B*_ = 0.441 for a Gaussian pulse shape. Transform limits reported for specific datasets in Sections 3 and 4 are instead computed directly from each measured spectrum, without assuming a Gaussian shape. We then applied the spectral phase expansion (Eq. (4)) to add varying amounts of GDD and TOD, obtained the time-domain field via inverse FFT, and calculated the third-order interferometric autocorrelation (Eq. (8)).

Fig. 2 shows four representative results. A transform-limited pulse with no added dispersion produces a narrow, symmetric THG-IAC trace, the shortest achievable for the given spectral bandwidth (Fig. 2a). Adding GDD (1700 fs^2^) stretches the pulse and broadens the envelope (Fig. 2b); adding TOD (42,000 fs^3^) instead redistributes energy into temporal side lobes (Fig. 2c). With both present, the trace reflects features of each – broadening from GDD and side lobes from TOD (Fig. 2d).

**Figure 2.**
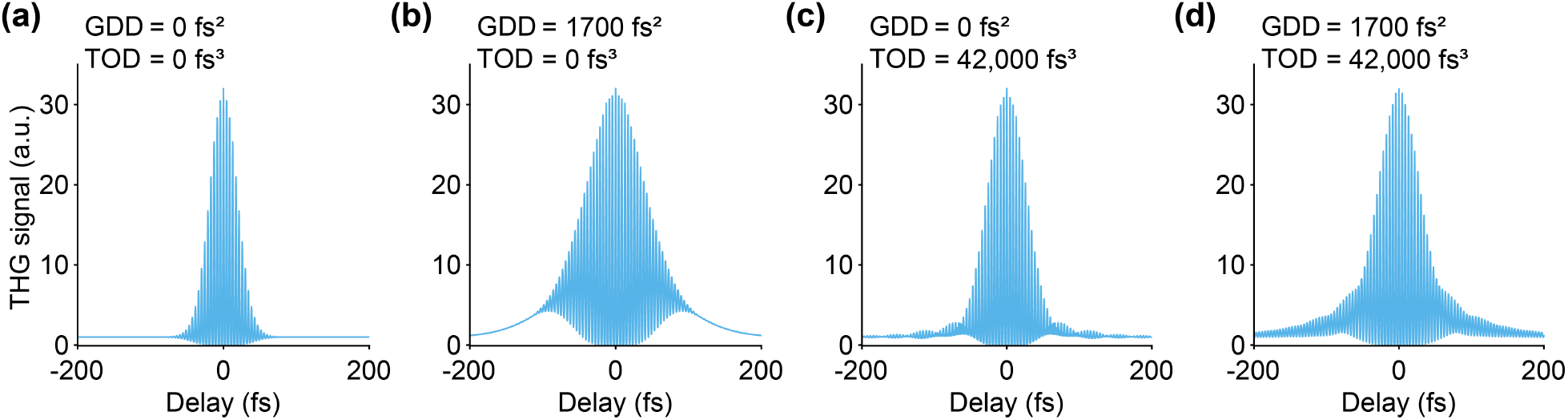
Simulated THG interferometric autocorrelation traces for Gaussian pulses at the 1300 nm window with varying amounts of dispersion. **(a)** Transform-limited pulse with no added GDD or TOD; a pulse duration of 34.8 fs was assumed, based on a spectral width of 69.6 nm centered at 1284.4 nm. **(b)** GDD (1700 fs^2^) stretches and chirps the pulse, broadening the THG-IAC trace. **(c)** TOD (42,000 fs^3^) redistributes energy into temporal side lobes. **(d)** Combined GDD and TOD produce both broadening and side lobes.

These simulations provide a visual reference for identifying dominant dispersion effects in measured THG-IAC traces and help guide compressor tuning during in situ pulse optimization.

### 2.5 Autocorrelation acquisition and data processing

THG-IAC traces, *G*^(3)^(*τ*), were obtained by recording the THG signal as a function of the temporal delay between two pulse replicas produced by the autocorrelation module. Data acquisition (vDAQ, MBF Bioscience) and motorized stage control were implemented with custom MATLAB code, which triggered constant-velocity stage motion and data collection. Details are given in Supplementary Note S3.

The laser pulses were focused on a water-glass interface to generate a strong THG signal. To eliminate spatial dependence, galvo mirrors were held stationary at the center of the field of view. The translation range was set to at least double the expected pulse overlap window to ensure the full THG-IAC trace was captured. Typical scans under 1300 nm excitation used a 0.14 mm stage translation (0.28 mm round-trip optical path length), corresponding to a delay range of approximately 1 ps.

Signal acquisition was synchronized to the laser pulses, with one sample collected per pulse within an 8 ns window at a fixed delay from the pulse trigger. After acquisition, delay values were computed from the recorded stage position, converted to round-trip time via the speed of light, giving a sample spacing of ∼6.67×10^-4^ fs. Each raw THG-IAC trace was smoothed using a moving-average filter of 400 samples, corresponding to a temporal window of approximately 0.27 fs (∼6% of the optical fringe period at 1300 nm and ∼5% at 1600 nm). The filter attenuates high-frequency fluctuations while preserving the interferometric fringe structure. The dark background – signal recorded on the same detector with no laser light – was averaged and subtracted from the smoothed trace, and the result was normalized to the mean of the first 10,000 datapoints (a low-signal region at the beginning of each acquisition, representing less than 1% of the total trace length). To identify τ = 0, a Gaussian fit was applied to a smoothed version of the trace and used to align paired 0-rod and 1-rod acquisitions with each other and with the simulated traces used in the fit. The centered trace was then linearly interpolated onto a uniform 0.2 fs delay grid within a ± 1100 fs window used by the fitting procedure (Section 2.6).

### 2.6 Dispersion estimation

To estimate the baseline group-delay dispersion (GDD_0_) and third-order dispersion (TOD_0_) of the excitation pulse at the sample plane, we developed a combined dispersion-estimation approach consisting of a look-up-table-based initialization step, which we call Dispersion Look-Up-Table Estimation (D-LUTE), followed by a joint two-measurement nonlinear fit. The forward model underlying this approach is described in Section 2.6.1. D-LUTE (Section 2.6.2) identifies candidate (GDD_0_, TOD_0_) regions from how the THG-IAC width changes as known dispersive glass rods are added to the beam path, and these candidate values initialize the grid search center used in the joint fitting procedure (Section 2.6.3). This procedure simultaneously fits two experimentally measured THG-IAC traces – one with no dispersive material in the beam path (0-rod) and one with a single 20 mm N-SF11 glass rod added (1-rod) – to a common forward model, with the N-SF11 rod’s known GDD and TOD increment (GDD_N-SF11_, TOD_N-SF11_) held fixed. This joint fit both reduces the sign ambiguity inherent to a single THG-IAC trace (Section 2.6.4) and provides a reference against which (GDD_0_, TOD_0_) are jointly determined. This method is validated on synthetic data with known ground truth in Section 3.1.

#### 2.6.1 Forward model

For a given pulse spectral amplitude *S*(*ω*), a simulated THG-IAC trace *G*^(3)^_sim_(*τ*; GDD_0_, TOD_0_) was computed by evaluating Eq. (5) with *φ*_2_ = GDD_0_ and *φ*_3_ = TOD_0_, inverse Fourier transforming to obtain *E*(*t*), and computing the third-order interferometric autocorrelation (Eq. (8)). *S*(ω) was Gaussian, with parameters from the fit to the measured spectrum. The forward model assumes the pulse spectral phase is fully described by GDD and TOD; higher-order or non-polynomial spectral phase, if present in the actual pulse, is not captured and would contribute additional error (see Section 4). A fourth-order term can be fit to the measurement pair to improve the fit agreement, although we cannot confirm whether it reflects true fourth-order dispersion (Supplementary Note S4).

#### 2.6.2 D-LUTE initialization

D-LUTE was used to generate initial dispersion (GDD_0_, TOD_0_) estimates for the joint fitting procedure. For the measured pulse spectrum, we precomputed a look-up table relating a range of simulated dispersion values to a measurable pulse observable: the width of the intensity autocorrelation, obtained from the interferometric autocorrelation via spectral filtering. By comparing how the measured 0-rod and 1-rod autocorrelation widths change against the known dispersion added by the N-SF11 rod, D-LUTE uses the look-up table to build a likelihood surface over candidate dispersion values. Anchoring the estimate to this differential change allows D-LUTE to suppress the non-trivial ambiguities that otherwise arise from pulse reconstruction via autocorrelation^9^, while preserving the experimentally simple, autocorrelation-based workflow. D-LUTE does not itself perform a complete pulse-field retrieval; it provides physically informed initial estimates for the subsequent joint fit. Full implementation details are given in Supplementary Note S5.

#### 2.6.3 Joint two-rod fitting procedure

For each rod condition (*m*= 0, 1), the simulated trace was evaluated at the total dispersion (GDD_0_ + *m*× GDD_N−SF11_, TOD_0_ + *m*× TOD_N−SF11_) and compared to experiment via an envelope-based cost function:

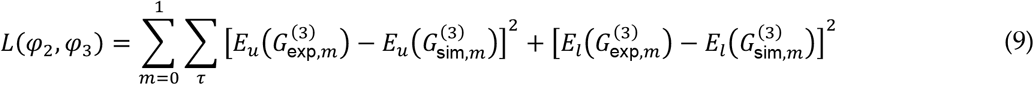

where *E*_*u*_ and *E*_*l*_ are the upper and lower envelopes, extracted identically for experimental and simulated traces. We used the envelope metric rather than raw fringe residuals because fringe positions are highly sensitive to sub-fringe delay drift between measurements, whereas the envelope shape encodes dispersion without requiring exact fringe alignment.

Each D-LUTE candidate was evaluated against the measured traces using Eq. (9), computed on envelopes spectrally filtered to isolate the 2*ω*_0_ component, which we found gave more noise-robust candidate selection; the candidate with the lowest envelope cost was carried forward as the starting point for the fit. The fit proceeded in two stages: a constrained grid search over (GDD_0_, TOD_0_) identified the global minimum in *L* within physically reasonable bounds, followed by nonlinear least-squares refinement (lsqnonlin, MATLAB) initialized at the grid minimum, using the envelope residuals as the fitting vector. If the refined solution fell near or beyond a grid boundary, the grid was recentered and the search repeated, allowing a compact grid to be used without restricting the accessible parameter range. The fitted baseline values (GDD_0_, TOD_0_) include the autocorrelator module’s dispersion and are reported in this form throughout the results. The corresponding residual dispersion during standard imaging is estimated by subtracting the BK7 beamsplitter’s calculated double-pass GDD and TOD contributions, given in Section 2.2. Full implementation details are given in Supplementary Note S6.

#### 2.6.4 Constraining pulse ambiguities

Autocorrelations do not uniquely define the pulses that create them. Pulses of different spectra or dispersion values may produce identical autocorrelation traces^9^. However, these traces do not all change in the same way when additional dispersion is added to the system. By focusing on how the autocorrelation width changes with added dispersion, D-LUTE favors the true dispersion values. Degenerate candidates that mimic the pulse’s autocorrelation no longer do so once significant known glass dispersion is applied.

The sign ambiguity is one instance of this: a single THG-IAC trace admits (GDD_0_, TOD_0_) and (−GDD_0_, −TOD_0_) as identical traces. Adding a dispersive rod can break this symmetry, as oppositely signed candidates predict different changes to the THG-IAC traces and consequently have different joint-fit costs. For the N-SF11 rods used here, the cost differences between oppositely signed GDD_0_ candidates are typically large and reliably identify the correctly signed candidate. By contrast, the cost differences between oppositely signed TOD_0_ candidates are small, indicating the added amount of TOD does not significantly change the traces. The joint-fit favors the positive-signed TOD_0_ candidate, but, given this small cost difference, may not robustly identify it in the presence of noise. We therefore additionally restricted the search to TOD_0_ > 0, consistent with the positive residual TOD reported for similar laser sources^5^. As shown in Supplementary Note S7, the joint-measurement cost function favors positive TOD even when this constraint is removed. The TOD_0_ > 0 constraint reflects both this cost-function preference and the literature precedent above. This ambiguity is demonstrated on synthetic data in Results, Section 3.1.

#### 2.6.5 Measurement uncertainty estimation

To assess measurement uncertainty, three independent THG-IAC traces were acquired per rod condition. The joint fit was repeated across all *N* = 3 × 3 = 9 permutations of (0-rod, 1-rod) trace pairs, and the mean and standard deviation of the recovered (GDD_0_, TOD_0_) across permutations were reported. This spread reflects experimental non-idealities not captured by the idealized noise analysis of Results, Section 3.1 – in particular, residual trace asymmetry from beam misalignment, which is present to some degree in every measurement and varies somewhat from trace to trace, perturbing the recovered dispersion more strongly than measurement noise alone. The resulting uncertainty therefore provides a more realistic estimate of the method’s precision under experimental conditions than the noise-limited characterization.

### 2.7 Mouse chronic cranial window implantation for in vivo brain imaging

All animal procedures were approved by the Yale University Institutional Animal Care and Use Committee. C57BL/6J mice (The Jackson Laboratory) were used for the experiments presented here. For in vivo mouse brain imaging, a chronic cranial window implantation procedure was performed. Briefly, mice were anesthetized with isoflurane (1-2% in O_2_) and administered carprofen (5 mg kg^−1^, subcutaneous) and lidocaine (2 mg kg^−1^, intradermal). Under aseptic conditions, a 5-mm-diameter craniotomy was created over the primary somatosensory cortex (S1), leaving the dura intact. The cranial window consisted of a donut-shaped coverslip (inner diameter 4.5 mm, outer diameter 5.5 mm; Potomac Photonics) bonded to a 5-mm-diameter coverslip (No. 1 thickness; Harvard Apparatus) using a UV-curable optical adhesive (Norland Optical Adhesive 61). The window was placed into the craniotomy and secured with cyanoacrylate glue. A titanium head post was attached to the skull using C&B Metabond (Parkell). Imaging was performed at least one week after surgery. During imaging, mice were head-fixed and maintained under isoflurane anesthesia (1-2% in O_2_).

## 3 RESULTS

### 3.1 Validation of dispersion recovery on synthetic data

Using the forward model and joint fitting procedure described in Section 2.6, we first validated recovery accuracy on synthetic 1300 nm data with known ground truth. For this validation the grid search was centered on the ground-truth values rather than initialized by D-LUTE, isolating the accuracy of the joint fit itself, independent of the initialization step used for experimental data. The grid spanned ±700 fs^2^ in GDD_0_ and ±30,000 fs^3^ in TOD_0_ about its center. Grid ranges used for the subsequent Sections on experimental fits were initialized by D-LUTE and shifted if the refined solution approached a grid boundary – enabling more constrained, efficient grid searches (Supplementary Note S6).

Fig. 3a,b (top, middle rows) show representative simulated 0-rod and 1-rod THG-IAC traces with added

**Figure 3.**
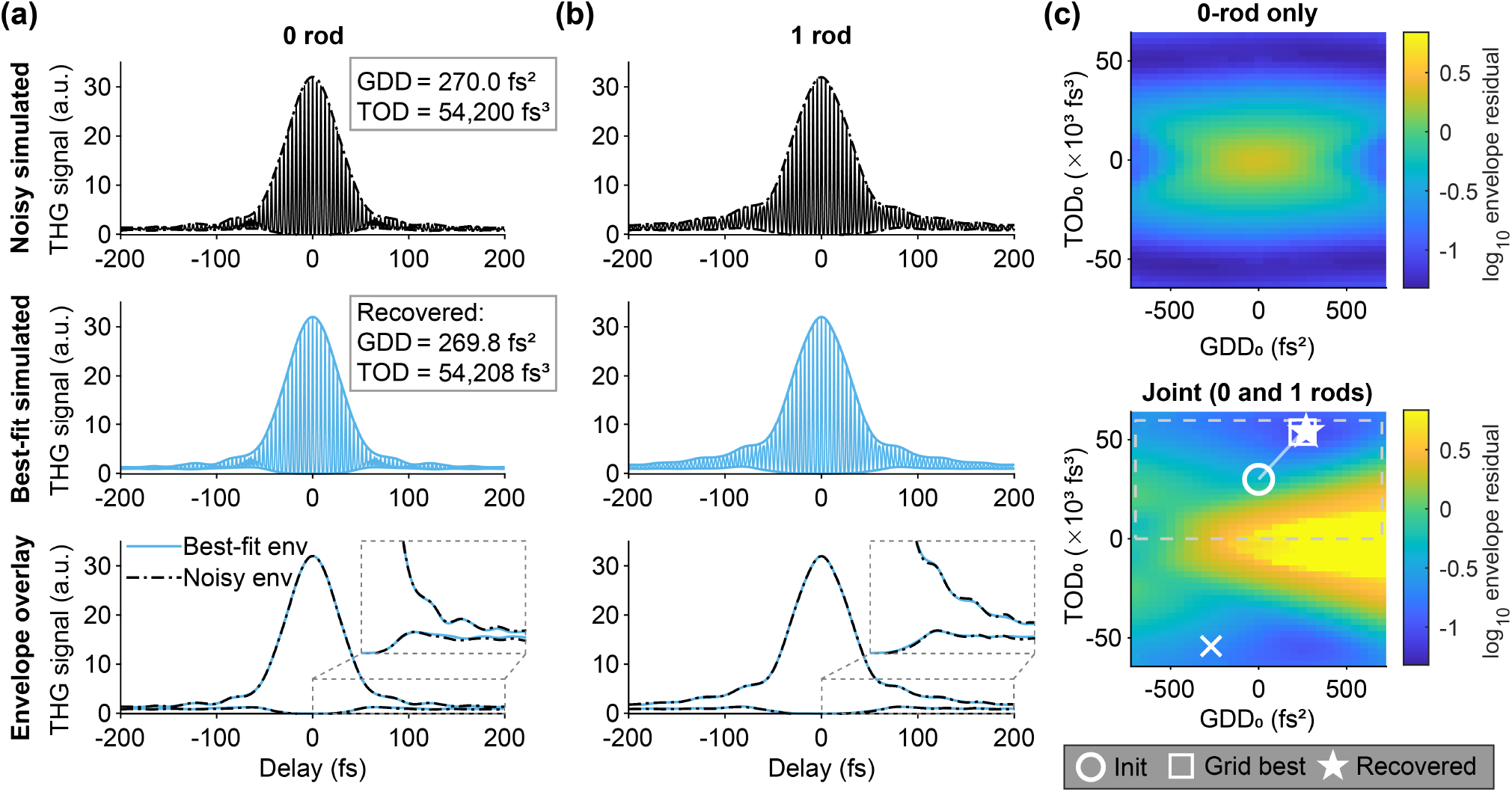
Validation of baseline-dispersion recovery on synthetic THG interferometric autocorrelation traces. Simulated 1300 nm THG-IAC traces for **(a)** 0-rod and **(b)** 1-rod configurations at baseline GDD_0_ = 270 fs^2^, TOD_0_ = 54,200 fs^3^ (recovered: 269.8 fs^2^, 54,208 fs^3^). (Top) Noisy simulated traces mimicking a measurement (noise level σ = 0.08), with extracted envelope. (Middle) Best-fit simulated traces at the recovered dispersion, with envelope. (Bottom) Overlay of noisy (dash-dot) and best-fit (solid) envelopes; the inset zooms in on the trailing wing. **(c)** Envelope-residual landscape over (GDD_0_, TOD_0_), log_10_ scale, for the 0-rod trace alone (top) and jointly fit with the 1-rod trace (bottom). The 0-rod residual shows two comparably low regions at positive and negative TOD_0_, reflecting the ambiguity of a single THG-IAC trace; adding the 1-rod trace raises the negative-TOD_0_ minima (×) above the true solution. Dashed box marks the search window (TOD_0_ > 0); markers show the recovery pipeline: initial guess (circle) → grid-search best node (square) → refined solution (star).

Gaussian noise (σ = 0.08, matching the measured experimental noise level from the trace background wings, |delay| > 250 fs), alongside the corresponding best-fit simulated trace at the recovered dispersion values. Fig. 3a,b (bottom row) show the best-fit envelopes overlaying the input envelopes across all delays, including the low-signal wings that carry most of the TOD information. Both configurations are fit jointly with a single (GDD_0_, TOD_0_); the 1-rod traces include the added dispersion of one 20 mm N-SF11 rod.

Fig. 3c demonstrates the sign ambiguity described in Section 2.6.4 and how joint fitting reduces it. Using the 0-rod THG-IAC trace alone, the residual forms two comparably low-cost regions at positive and negative TOD_0_ (top), so neither the sign of TOD_0_ nor the value of GDD_0_ is determined. Jointly fitting the 0- and 1-rod traces reduces this degeneracy (bottom): the true solution becomes the global minimum, while the negative-TOD_0_ minima are raised above it, though by a modest margin. Markers trace the recovery pipeline from initial guess through grid-search minimum to refined solution.

To characterize recovery accuracy, we generated synthetic 0-rod and 1-rod THG-IAC traces across a grid of baseline (GDD_0_, TOD_0_) values, added Gaussian noise of standard deviation σ, and applied the recovery procedure, averaging the bias over multiple noise realizations at each grid point. GDD_0_ bias is reported in absolute units (fs^2^) because GDD_0_ spans zero, where a fractional bias would diverge; TOD_0_ bias is reported as a percentage, being bounded away from zero. Fig. 4a shows the resulting bias maps at the estimated noise level (σ = 0.08, mean over 20 noise realizations); recovery is accurate across the plane, with mean GDD_0_ bias of a few tens of fs^2^ and mean TOD_0_ bias of a few percent.

**Figure 4.**
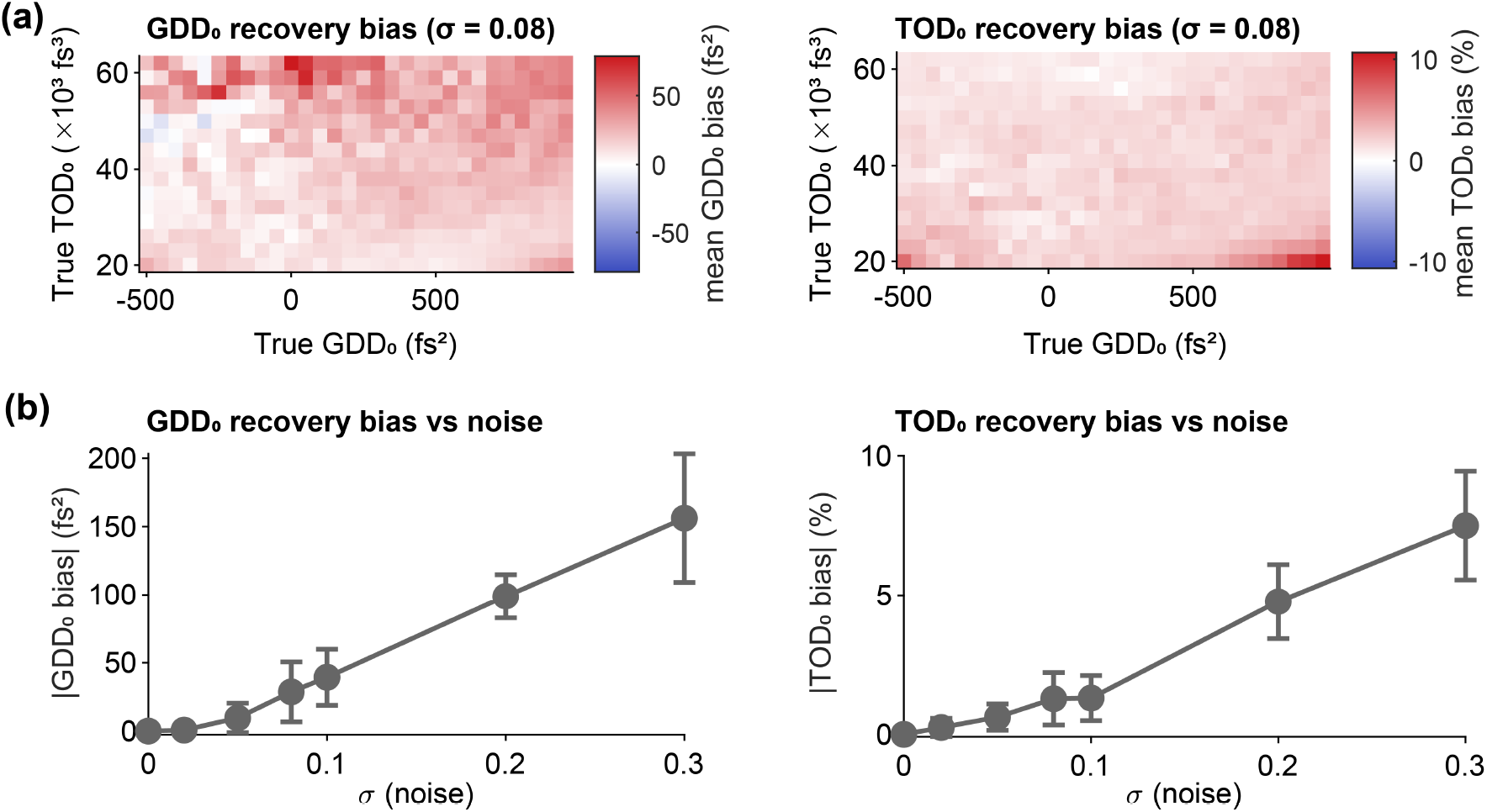
Recovery accuracy and precision of baseline dispersion from synthetic data. **(a)** Recovery bias across the plane of true (GDD_0_, TOD_0_) at the estimated noise level (σ = 0.08, mean over 20 noise realizations): (left) GDD_0_ bias (fs^2^) and (right) TOD_0_ bias (%). **(b)** Recovery bias versus noise level σ at a representative operating point (GDD_0_ = 270 fs^2^, TOD_0_ = 54,200 fs^3^); markers show the absolute mean bias and error bars the standard deviation over 30 noise realizations: (left) |GDD_0_ bias| and (right) |TOD_0_ bias|.

Fig. 4b shows the recovery bias and its standard deviation across noise realizations versus noise level σ at a representative operating point; both GDD_0_ and TOD_0_ biases remain small across the range and grow gradually with noise.

These values reflect the bias and noise-limited scatter under the idealized assumption that measurement noise is the only error source, and the forward model exactly describes the data. In practice, additional error arises from experimental non-idealities – such as residual THG-IAC trace asymmetry from beam misalignment (Section 2.6.5) – and from departures of the true pulse spectral phase from the assumed GDD/TOD model (see Section 4), either of which can exceed this idealized bias-and-scatter estimate.

### 3.2 THG interferometric autocorrelation measurements at 1300 nm

THG-IAC traces were acquired at 1300 nm with no dispersive material in the beam path (0-rod) and with a single 20 mm N-SF11 glass rod inserted (1-rod). To identify the compressor setting giving the smallest GDD at the focal plane, traces were acquired across a range of compressor settings, with three independent THG-IAC traces acquired per rod condition at each setting. Although the method requires only a single trace per rod condition, the joint two-rod fitting procedure (Section 2.6.3) was applied across all N = 3 × 3 = 9 trace-pair permutations at each setting, recovering the baseline dispersion (GDD_0_, TOD_0_) and estimating its uncertainty (Section 2.6.5). This identified −5400 fs^2^ (Supplementary Note S8 and Fig. S7) as the optimal setting, within the resolution of the settings sampled; all results below correspond to it.

Fig. 5a,b show a representative 0-rod and 1-rod trace pair measured at the optimal compressor setting on this system (Laser 1), together with the corresponding best-fit simulated traces and envelope overlays used to assess fit quality. The fit closely reproduces the upper and lower envelope shape across the full delay range. Across the 9 permutations, the recovered baseline dispersion was GDD_0_ = 289 ± 99 fs^2^ and TOD_0_ = 54,400 ± 4,900 fs^3^ (mean ± std, Fig. 5c). Applying the forward model to each of the 9 recovered (GDD_0_, TOD_0_) pairs yielded a baseline pulse duration of *τ*_*p*_= 55 ± 1 fs (mean ± std). For comparison, the transform-limited pulse duration, obtained by inverse Fourier transforming the measured spectral amplitude *S*(*ω*) assuming zero spectral phase, is 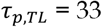 fs.

**Figure 5.**
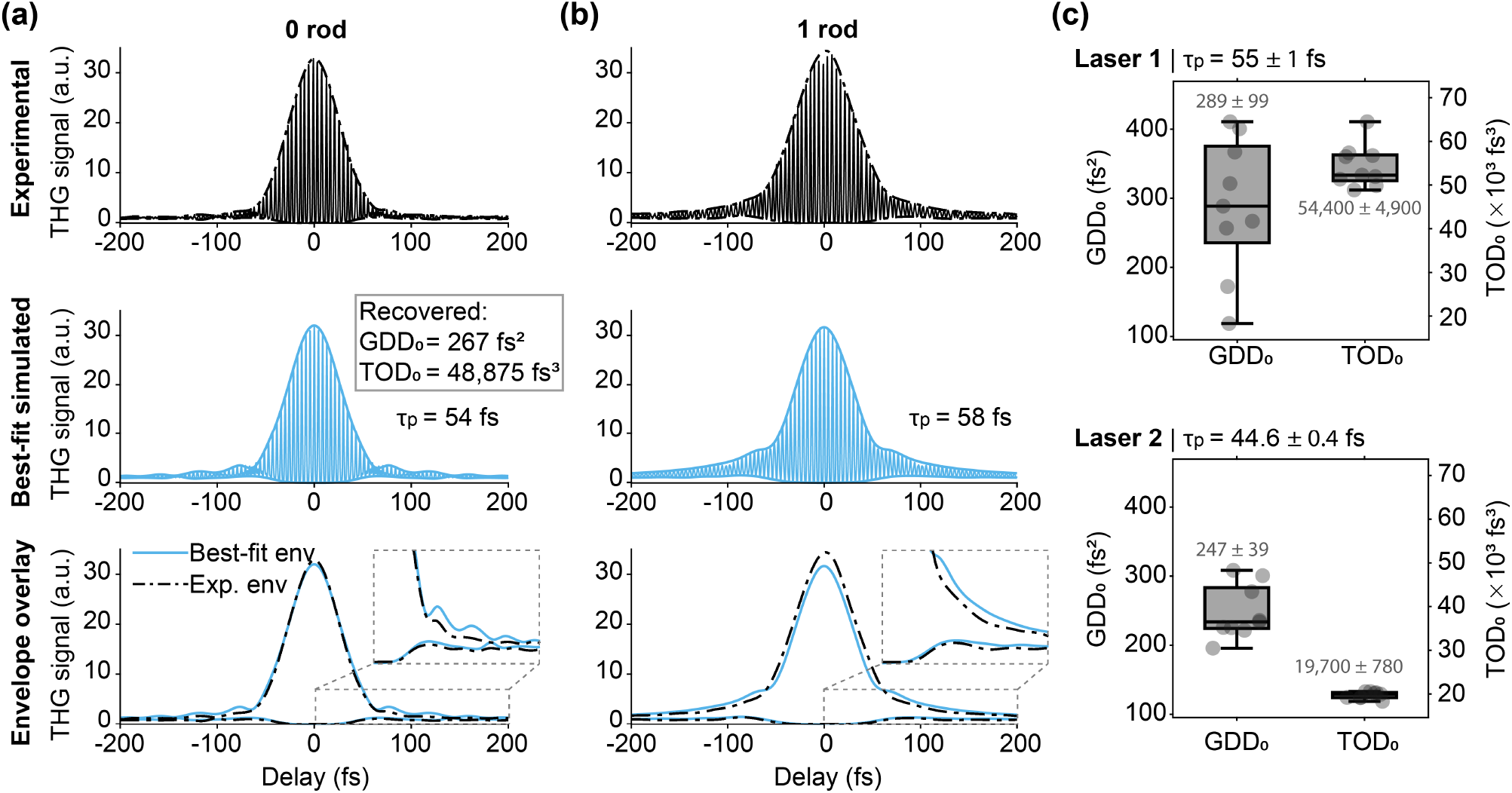
Dispersion estimation from THG interferometric autocorrelation measurements at 1300 nm. **(a,b)** Representative 0-rod **(a)** and 1-rod **(b)** experimental THG-IAC traces at the optimal compressor setting on Laser 1, with best-fit simulated traces and envelope overlays; GDD_0_ and TOD_0_ were recovered by jointly fitting the paired 0-rod/1-rod measurements, with their relative dispersion constrained by the known dispersion of the N-SF11 rod. For this representative pair, the recovered baseline dispersion was GDD_0_ = 267 fs^2^, TOD_0_ = 48,875 fs^3^, corresponding to a pulse duration of *τ*_*p*_ = 54 fs (0-rod) and 58 fs (1-rod). **(c)** Across N = 9 permutations, the recovered baseline dispersion was GDD_0_ = 289 ± 99 fs^2^ and TOD_0_ = 54,400 ± 4,900 fs^3^ (mean ± std) on Laser 1, and GDD_0_ = 247 ± 39 fs^2^ and TOD_0_ = 19,700 ± 780 fs^3^ on Laser 2 (same laser model), at their respective optimal settings.

To test the generality and practicality of the method, the autocorrelator module was installed on a second microscope built with matching components (Section 2.1), equipped with a different unit of the same laser model (Cronus 3P, Light Conversion; Laser 2). The same procedure yielded GDD_0_ = 247 ± 39 fs^2^ and TOD_0_ = 19,700 ± 780 fs^3^ at its optimal setting (Fig. 5c), corresponding to a mean baseline pulse duration of *τ*_*p*_ = 44.6 ± 0.4 fs (computed as above), compared to a transform limit of 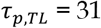fs.

While GDD_0_ was comparable between the two systems, TOD_0_ differed by a factor of approximately 2.7 despite the identical laser model. Both lasers’ characteristics, which we monitor regularly, remained stable across days, shutdowns, and restarts, indicating that the TOD discrepancy between the two lasers is not attributable to short-term instability.

### 3.3 THG interferometric autocorrelation measurements at 1600 nm

Longer excitation wavelengths improve tissue penetration for both 2-photon^28^ and 3-photon microscopy^1^. To characterize pulse dispersion in this regime, we applied our autocorrelator module at 1600 nm excitation, where several glasses and crystals in the microscope beam path exhibit anomalous dispersion, producing negative GDD at the sample plane. In our system, this is compensated by tuning the internal prism compressor to introduce positive GDD, offsetting the microscope’s intrinsic negative dispersion.

At 1600 nm, N-SF11 contributes substantially less GDD per unit length than at 1300 nm, so a single rod produces only a small, difficult-to-resolve shift in dispersion. We therefore used 0-rod and 3-rod conditions to obtain a larger, more reliably fit dispersion increment. As at 1300 nm, three independent THG-IAC traces were acquired per rod condition across a range of compressor settings, and the joint fitting procedure (Section 2.6.3) was applied across all N = 3 × 3 = 9 permutations; all results below correspond to the 4500 fs^2^ compensation setting (Supplementary Note S8 and Fig. S8).

Fig. 6a,b show a representative 0-rod and 3-rod THG-IAC trace pair at the 4500 fs^2^ compensation setting, with the corresponding best-fit simulated traces and envelope overlays. The recovered baseline dispersion across the 9 permutations was GDD_0_ = −1,140 ± 320 fs^2^ and TOD_0_ = 118,200 ± 3,500 fs^3^ (mean ± std, Fig. 6c), corresponding to a mean baseline pulse duration of *τ*_*p*_ = 77 ± 1 fs (mean ± std across 9 permutations, as in Section 3.2), compared to a transform limit of 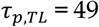 fs. This compressor setting was chosen as representative because it corresponds to a residual GDD near zero at the sample plane during standard imaging, once the BK7 beamsplitter’s own double-pass contribution (present only while the autocorrelator module is in the beam path) is taken into account. Setting GDD_0_ to zero in the forward model while holding TOD_0_ fixed shortens the calculated pulse duration by only ∼1 fs, indicating that TOD, not residual GDD, is what keeps the pulse from reaching its transform limit.

**Figure 6.**
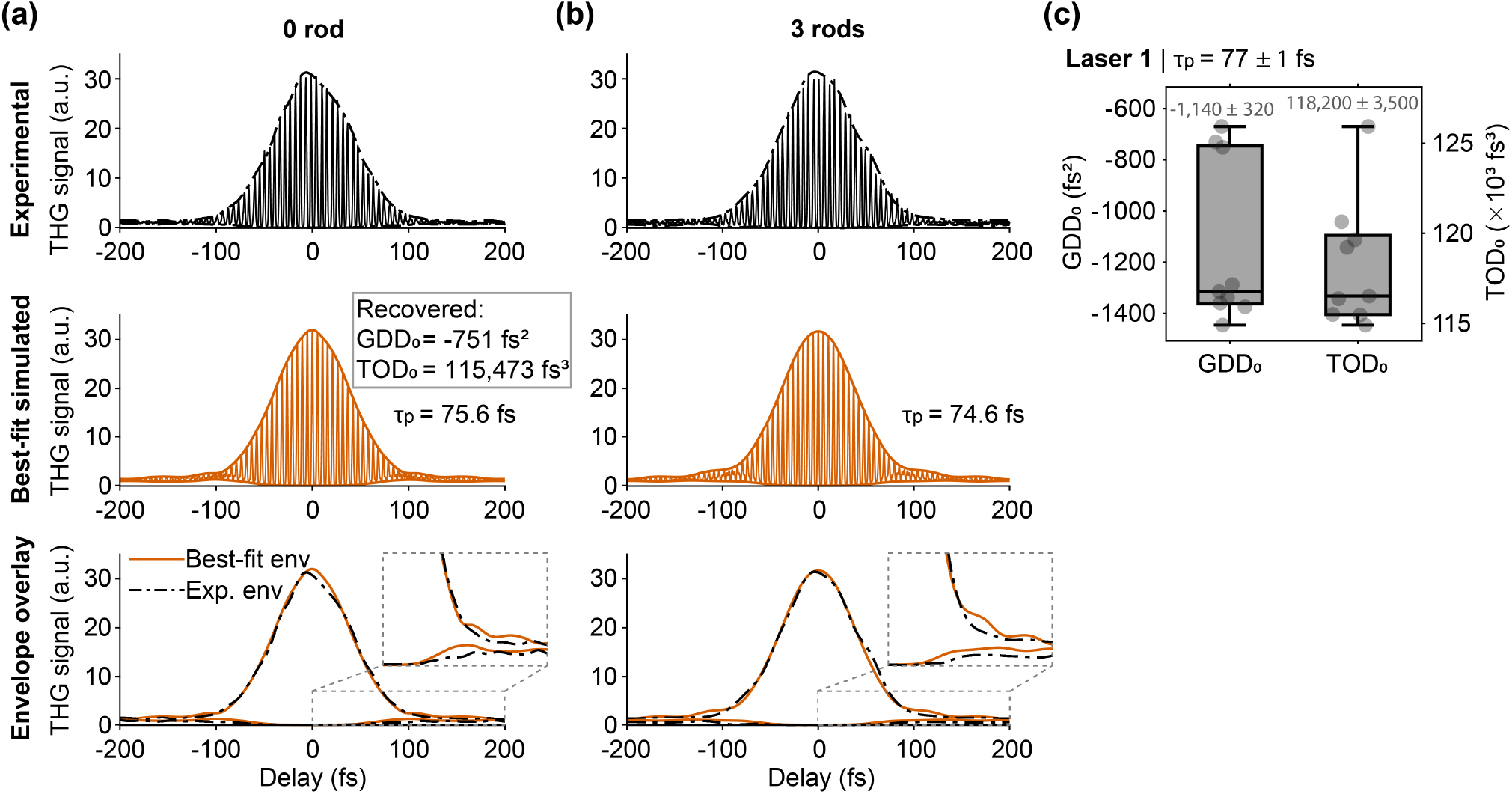
Dispersion estimation from THG interferometric autocorrelation measurements at 1600 nm. **(a,b)** Representative 0-rod **(a)** and 3-rod **(b)** experimental THG-IAC traces at the optimal compressor setting for standard imaging (4500 fs^2^ compensation), with best-fit simulated traces and envelope overlays; GDD_0_ and TOD_0_ were recovered by jointly fitting the paired 0-rod/3-rod measurements, with their relative dispersion constrained by the known dispersion of the N-SF11 rods. For this representative pair, the recovered baseline dispersion was GDD_0_ = −751 fs^2^, TOD_0_ = 115,473 fs^3^, corresponding to a pulse duration of *τ*_*p*_ = 75.6 fs (0-rod) and 74.6 fs (3-rod). **(c)** Across N = 9 permutations, the recovered baseline dispersion was GDD_0_ = −1,140 ± 320 fs^2^ and TOD_0_ = 118,200 ± 3,500 fs^3^ (mean ± std).

### 3.4 In vivo pulse characterization deep in mouse brain tissue

A key advantage of using THG as the nonlinear signal source for interferometric autocorrelation is that, unlike second-order autocorrelators that require a dedicated nonlinear crystal, THG can be generated directly by many endogenous biological structures^21^. In the brain, myelin sheaths surrounding axons are a particularly strong source of THG contrast, owing to their lipid-rich, layered membrane structure^29,30^. This allows pulse characterization to be performed directly within intact tissue, without introducing an artificial interface.

These structures were used to measure pulse dispersion in vivo at 1300 nm excitation (Laser 2) through a cranial window in the mouse brain (Fig. 7). We first acquired a THG-IAC trace at the glass-brain interface, just below the cranial window (Fig. 7a), providing a reference measurement near the tissue surface. We then translated the focus to a depth of approximately 900 µm below the brain surface (i.e., bottom of the cranial window), into white matter, and used the endogenous THG signal from myelinated fibers (Fig. 7b) to acquire a second THG-IAC trace at depth. The two THG-IAC traces overlap closely (Fig. 7c): the FWHM of the traces at the two locations were compared using a two-tailed Welch’s t-test (N = 5 traces per location), revealing no statistically significant difference (surface: 55.9 ± 0.6 fs; depth: 57 ± 2 fs; p = 0.18), consistent with negligible additional dispersion at the focus between the tissue surface and a depth of nearly a millimeter. This result demonstrates that our autocorrelator module can be used to directly verify pulse quality at depth in biological tissue in vivo. Although no measurable additional dispersion was found at the focal plane under these conditions, other tissues may behave differently, and this module provides a direct way to check.

**Figure 7.**
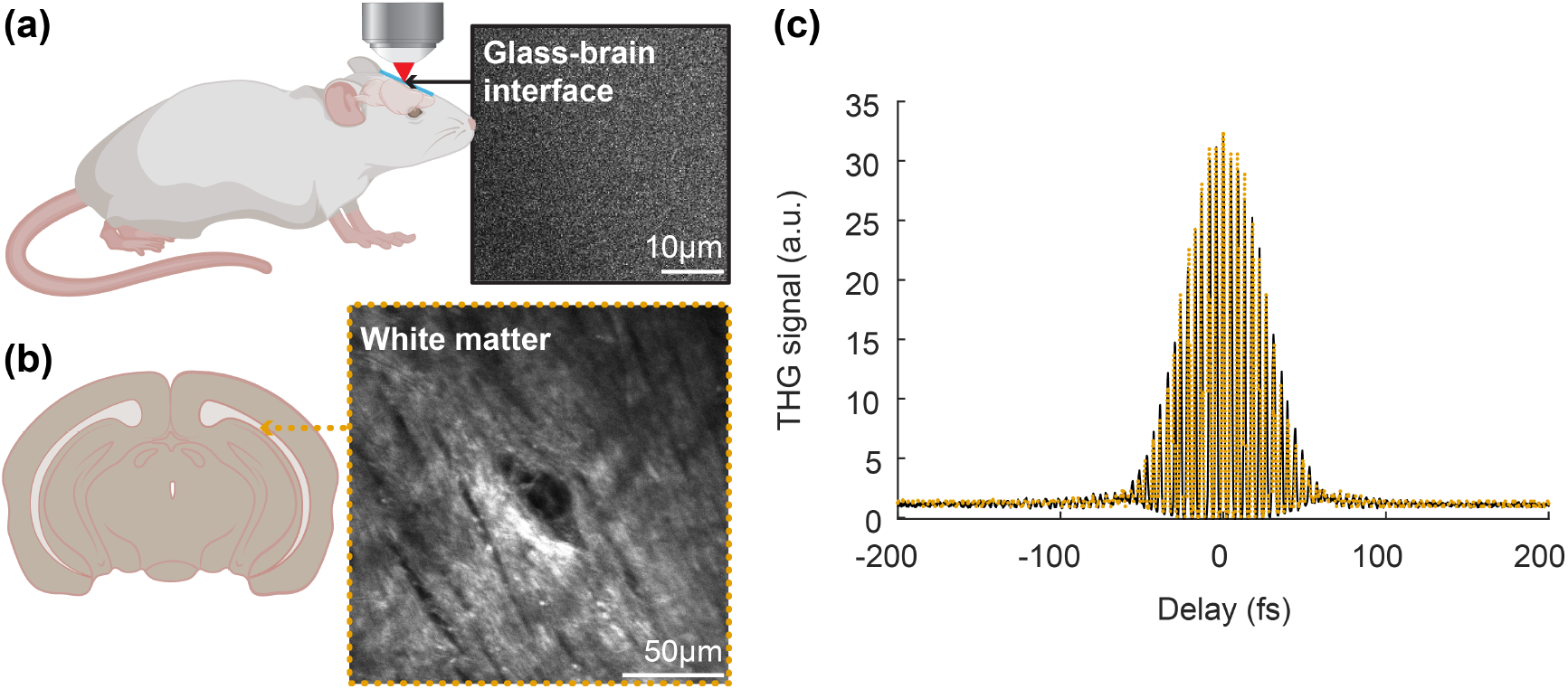
In vivo pulse characterization in the mouse brain. **(a)** Cranial window preparation for in vivo imaging (left) and representative THG image at the glass-brain interface, used to acquire a reference THG-IAC trace (right). **(b)** Coronal section of mouse brain indicating white matter layer imaged (left) and THG image of myelinated axons in white matter, ∼900 µm below the brain surface (i.e., bottom of the cranial window), used to acquire a second THG-IAC trace (right). **(c)** Overlay of traces acquired at the surface (black, solid) and in white matter (orange, dotted). FWHM did not differ significantly between surface (55.9 ± 0.6 fs) and depth (57 ± 2 fs; p = 0.18, Welch’s t-test, N = 5 each).

## 4 DISCUSSION

With the autocorrelator module in the beam path, the pulse duration measured at the focal plane greatly exceeded the transform limit at every condition studied here – 55 vs. 33 fs at 1300 nm (Laser 1), 44.6 vs. 31 fs at 1300 nm (Laser 2), and 77 vs. 49 fs at 1600 nm – a shortfall driven almost entirely by TOD. Because 3P excitation scales with the cube of instantaneous intensity, the cost is worse than the pulse duration alone suggests. With the signal scaling as 1/τ_p_^2^ at fixed pulse energy, this predicts a signal fraction (*τ*_*p,TL*_ /*τ*_p_)^2^ = 36% of the transform-limited value, whereas the full forward model applied to the recovered (GDD_0_, TOD_0_) values for Laser 1 at 1300 nm gives 22% (Supplementary Note S9 and Fig. S9). Most of this difference reflects the energy that TOD moves out of the central lobe and into the temporal wings (Fig. 2c). Setting GDD_0_ to zero while holding TOD_0_ at its recovered value raises the predicted signal marginally from 22% to 23% and shortens the predicted pulse from 55.0 fs to 54.7 fs. This indicates that residual TOD, not GDD, is the dominant barrier to reaching the transform limit.

This large TOD magnitude is not unique to a single unit: across the two nominally identical laser systems studied here, TOD_0_ differed by a factor of ∼2.7 despite comparable GDD_0_, showing that baseline dispersion cannot be assumed from the laser model alone and underscoring the value of a module that can measure it directly, in situ, with minimal setup. By combining D-LUTE initialization with a joint two-condition fit, this approach recovers both GDD_0_ and TOD_0_ while preserving the experimental simplicity of autocorrelation. The recovered TOD_0_ values are also consistent in magnitude with the only other reported measurement of baseline TOD_0_ for a similar 3PM laser source, made at the laser head rather than post-objective (∼22,750 fs^3^)^5^.

The laser’s built-in prism compressor, however, is designed to introduce and tune GDD rather than TOD, and so it cannot be used to jointly compensate both terms. Residual TOD of this magnitude could in principle be compensated using passive approaches such as grating pairs^31^ or grisms^32^. These simple approaches provide GDD and TOD at a fixed ratio set by their design, so they cannot correct the two independently, and that ratio is itself wavelength-dependent. More experimentally complex adaptive pulse-shaping strategies, such as those employing liquid-crystal spatial light modulators^33,34^, do offer independent and reconfigurable control of GDD and TOD – and, more generally, of arbitrary spectral phase, including higher-order and non-polynomial terms – providing a route toward transform-limited pulses and higher nonlinear signal at the focal plane. To estimate this benefit for standard imaging, we subtract the autocorrelator module’s BK7 beamsplitter contribution from the recovered baseline dispersion, giving GDD = 150 fs^2^ and TOD = 51,200 fs^3^, for Laser 1 at 1300 nm. Full compensation of both terms is then predicted to increase the signal ∼4.2-fold at fixed pulse energy, or equivalently reduce the excitation power by 38%, at equal signal.

Although the sign of TOD is not fully constrained by the measurement, our joint-fit cost function favors a positive value (Supplementary Note S7), and the modeled pulse duration and THG signal are, in any case, approximately symmetric with respect to the sign of TOD (Fig. S9; Supplementary Note S9). Thus, a negative TOD of the same magnitude would be expected to produce similar pulse broadening and THG signal, with only the sign of the required compensation differing.

The synthetic data characterization of Section 3.1 provides a noise-limited estimate of recovery accuracy and precision at the experimental noise level (σ ≈ 0.08): a recovery bias of a few tens of fs^2^ in GDD_0_ and a few percent in TOD_0_ (Fig. 4a), with a comparable scatter across noise realizations at a representative operating point (Fig. 4b). The experimental permutation spreads at 1300 nm (Fig. 5c) are comparable to or larger than this idealized scatter, consistent with the expectation (Section 2.6.5) that experimental non-idealities such as residual trace asymmetry, rather than measurement noise alone, set the achievable precision.

Separately from this, agreement between measured and best-fit simulated envelopes is good across all datasets shown in this study, but small residual discrepancies remain, concentrated in the THG-IAC trace wings where TOD’s signature is strongest. These differences may reflect limitations of the truncated polynomial spectral-phase model, experimental non-idealities in the measured traces, or both. The present measurements do not isolate their relative contributions. Eq. (4) truncates the spectral phase at third order and therefore cannot capture any fourth-order (FOD) term if present. Fixing GDD_0_ and TOD_0_ and fitting an additional FOD_0_ term does improve the envelope agreement (Supplementary Note S4), but we cannot confidently attribute this improvement to true fourth-order dispersion in the pulse rather than to residual trace asymmetry, measurement noise, an incorrectly identified carrier frequency, or non-polynomial phase that a finite-order polynomial expansion cannot represent – and that the added free parameter could instead be absorbing. Without an independent spectral-phase measurement (e.g., FROG or SPIDER) at the sample plane, any higher-order phase intrinsic to the OPA output remains unconstrained by this method.

Beyond 3PM at 1300 and 1600 nm, this module applies directly to other multiphoton modalities sharing the same THG-capable microscope and laser platform, provided the excitation wavelength is long enough for the resulting THG to fall within the transmission range of standard microscope optics – in practice, excitation above roughly 1150 nm. Within this range, the approach applies to THG imaging itself, 3PM, and even 2PM of far-red fluorophores^35,36^. In every case, the resulting THG signal could be monitored simultaneously using the same detection scheme already employed here, extending in situ pulse characterization across microscopy modalities and excitation wavelengths.

## 5 CONCLUSIONS

Using a third-order interferometric autocorrelation approach based on THG, together with D-LUTE initialization and a joint two-measurement fit, we estimated baseline GDD_0_ and TOD_0_ of femtosecond pulses directly at the focal plane of a multiphoton microscope under 1300 nm and 1600 nm excitation. This in situ approach requires only a compact autocorrelator module added to the beam path; the THG signal is collected on the microscope’s existing detection channel, with filters matched to the THG wavelength, rather than a separate detector or external instrument.

Measured pulse durations at the focal plane were 1.4-1.7× their transform limit at 1300 nm and 1600 nm, a shortfall driven almost entirely by TOD. For nonlinear signals with cubic intensity dependence (e.g., THG or 3PM), this residual TOD is predicted to reduce the achievable signal to roughly 22% of the value it would reach if the pulse were transform-limited, as estimated for Laser 1 with the autocorrelator module in the beam path (24% for the standard-imaging configuration). TOD_0_ itself varied by roughly 2.7-fold between nominally identical laser units, underscoring the need to measure it directly rather than assume it from the laser model.

In vivo, we used the endogenous THG signal from myelinated fibers in mouse brain to acquire interferometric autocorrelation traces at the tissue surface and at a depth of nearly a millimeter, finding no significant broadening between the two – demonstrating that pulse quality at depth can be verified directly using this approach.

Because THG can be efficiently generated and collected across the near-infrared excitation windows used in 3PM and longer-wavelength 2PM, this approach offers a practical, low-cost route to routine, in situ pulse monitoring across these nonlinear microscopy platforms. To support adoption, we provide the full analysis pipeline together with a standalone graphical user interface (GUI) that performs the dispersion estimation on user-supplied traces.

## ACKNOWLEDGMENTS

We thank Vasily Goncharov (SuperPostDoc LLC) for his support in the design and construction of our custom multiphoton microscopes, based on the Janelia MIMMS platform (Howard Hughes Medical Institute, Janelia Research Campus), and Axiom Optics for generously providing a demonstration d-vision spectrometer used for pulse spectral characterization during this work. We also thank Shengqi Wang for his contributions to building and troubleshooting the hardware and software of our custom multiphoton microscope.

## AUTHOR CONTRIBUTIONS

**Liam Shaughnessy:** Methodology (lead); Software (lead); Formal analysis (lead); Investigation (autocorrelator module construction and troubleshooting; implementation of the optimal data acquisition procedure); Writing – original draft (lead); Writing – review & editing.

**Lauren Vannell:** Investigation (experimental data acquisition; autocorrelator module construction and troubleshooting; implementation of the optimal data acquisition procedure); Formal analysis (in vivo data); Writing – original draft (supporting); Writing – review & editing.

**Sulekh Fernando-Peiris:** Investigation (cranial window surgeries; in vivo experiments); Supervision (in vivo experiments); Writing – review & editing.

**Cristina Rodríguez:** Conceptualization (lead); Project administration; Funding acquisition; Supervision (experiments and computational modeling); Methodology (supporting); Formal analysis (supporting); Writing – original draft (lead); Writing – review & editing (lead).

## CODE AND DATA AVAILABILITY

All MATLAB code used to simulate THG interferometric autocorrelation traces, analyze experimental data, and perform the joint dispersion-estimation fitting, together with sample datasets and documentation describing the repository structure and instructions for reproducing the figures and results presented in this manuscript, is available at https://github.com/rodriguezlabyale/thg-autocorrelator. A standalone GUI is also provided, allowing users to upload their own autocorrelation traces and perform the dispersion-estimation fitting directly.

