## Supplementary Information for "In situ pulse dispersion estimation via third-harmonic generation interferometric autocorrelation for multiphoton microscopy"

[1] Department of Applied Physics, Yale University, New Haven, CT, USA
[2] Department of Physics, Yale University, New Haven, CT, USA
[3] Department of Biomedical Engineering, Yale University, New Haven, CT, USA
[4] Wu Tsai Institute, Yale University, New Haven, CT, USA
[5] Howard Hughes Medical Institute, Chevy Chase, MD, USA
[†] These authors contributed equally to this work

### Supplementary Note S1: Autocorrelator module components

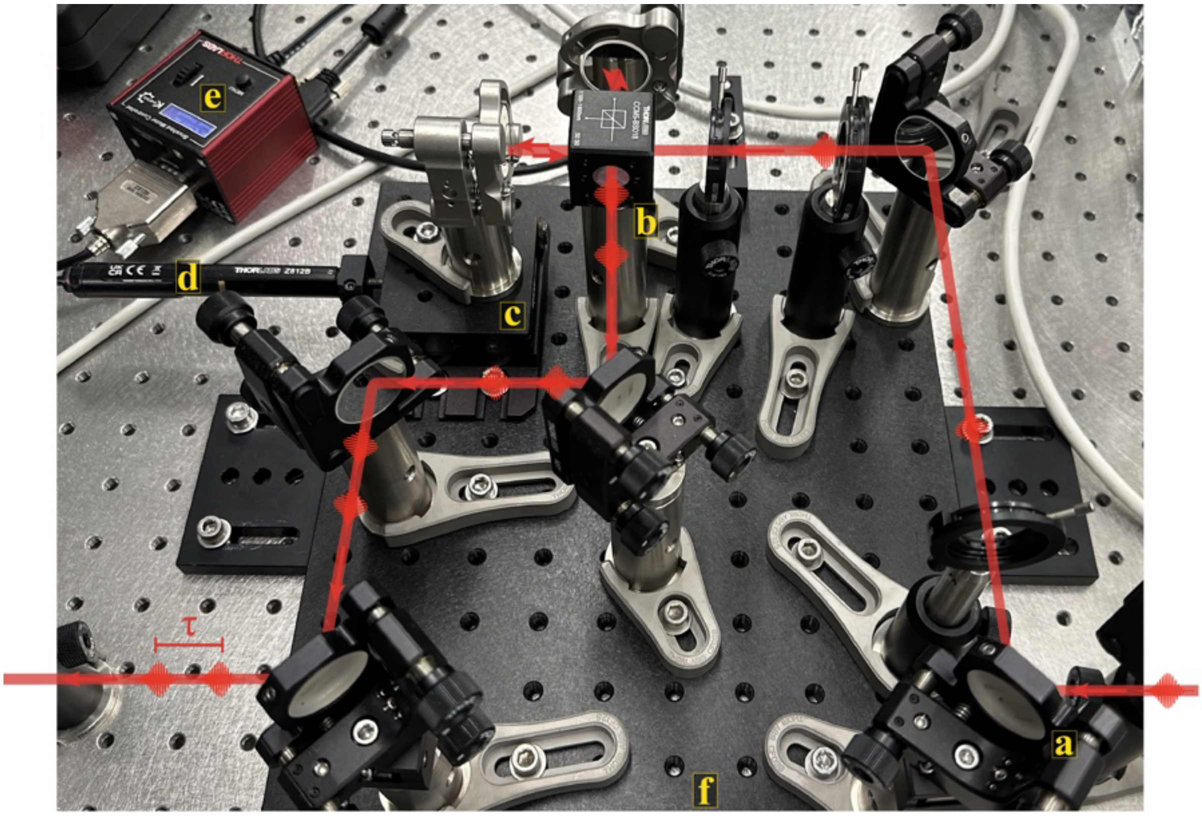


**Figure S1. | Autocorrelation module set-up and beam path.** First, two silver protected mirrors **(a)** align the excitation beam through the autocorrelation module. The 50:50 non-polarizing beamsplitter cube **(b)** separates the light into the two interferometric arms. One mirror of one arm is mounted on a translation stage **(c)** moved by a motorized actuator **(d)** connected to a K-Cube controller **(e)**, allowing for variable time delay (τ) between the two pulse replicas. The pulses are recombined at the beamsplitter and aligned back through the excitation beam path with two exit mirrors. The module is housed on a 10”x12” breadboard **(f)** for ease of movement.

**Table S1.** | **Autocorrelator module main components.**

| Item name | Cost (USD) | Amount |
|---|---|---|
| **a)** Silver protected mirror (PF10-03-P01) | 60.20 | 7 |
| **b)** 16 mm Cage Cube-Mounted Non-Polarizing Beamsplitter, 50:50 (R:T), 1100-1600 nm, 8-32 Tap (CCM5-BS018) | 290.00 | 1 |
| **c)** 1/2" Translation Stage with 1/4"-170 Adjuster, 1/4"-20 Taps (MT1B) | 278.36 | 1 |
| **d)** 12 mm Motorized Actuator, 3/8" Barrel Fitting (0.5 m cable) (Z812B) | 938.60 | 1 |
| **e)** K-Cube Brushed DC Servo Motor Controller (KDC101) | 826.75 | 1 |
| **f)** Aluminum Breadboard 10" x 12" x 1/2", 1/4"-20 Taps (MB1012) | 164.48 | 1 |
| **TOTAL** | **2,919.59** | |

**Table S2.** | **Additional optics.**

| Item name | Cost (USD) | Amount |
|---|---|---|
| Flip Mount Adapter, Imperial (FM90) | 102.19 | 2 |
| Ø1/2" Stainless Steel Optical Post - Imperial (TR3) | 6.58 | 8 |
| Standard Ø1/2" Post Holder (PH3) | 10.29 | 8 |
| Ø1.25" Studded Pedestal Base Adapter, 1/4"-20 Threads (BE1) | 11.59 | 8 |
| Clamping Forks (1.24”, captive screw) (CF125C) | 13.73 | 8 |
| Ø1" Precision Kinematic Mirror Mount, 3 Adjusters (KS1) | 106.54 | 7 |
| **TOTAL** | **1,287.68** | |

The maximum cost of this homemade system is $4,207.27, considerably less expensive than commercially available models, with room to further cut costs. All autocorrelator module components were purchased from Thorlabs Inc.

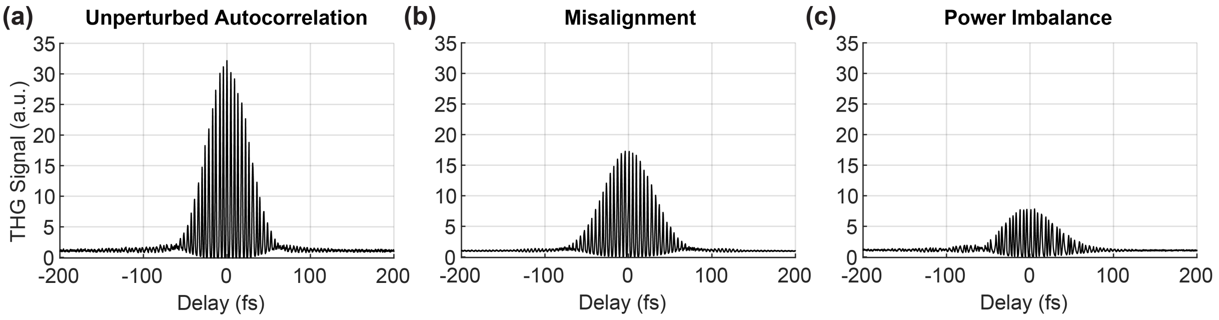


**Figure S2.** | **Representative problematic autocorrelation traces, with an ideal trace for comparison. (a)** An ideal autocorrelation trace with proper symmetry and peak-to-background ratio of 32:1. **(b)** An autocorrelation trace taken while both arms of the interferometer were not fully overlapped, with proper symmetry but a reduced peak-to-background ratio. **(c)** An autocorrelation trace taken with one arm of the interferometer decreased to 40% power, with an asymmetric envelope and reduced peak-to-background ratio.

### Supplementary Note S2: Spectral model selection for pulse simulations

The laser spectrum at 1300 nm was measured as described in Section 2.1 and is shown in Fig. S3. Gaussian and sech² fits were performed in the frequency domain by minimizing the normalized sum of squared residuals ($NSSR = \Sigma(y_{fit} - y_{data})^2 / \Sigma y_{data}{}^2$). The sech² model yielded a marginally lower NSSR (0.002999 vs. 0.003163, ratio 1.055), indicating a slightly better fit to the spectral shape, particularly in the wings. The measured spectrum exhibits slight asymmetry that is not captured by either symmetric model (Fig. S3).

To determine which model best represents the pulse for simulation purposes, we compared simulation outputs using three spectral inputs: the experimental spectrum directly, the Gaussian fit, and the sech² fit. For each, the pulse was represented in the frequency domain (with flat spectral phase) and transformed to the time domain via inverse FFT, and the third-order interferometric autocorrelation was computed. Table S3 summarizes the results.

The Gaussian model closely reproduces the results obtained using the experimental spectrum across all metrics. The 2nd order intensity autocorrelation FWHM to pulse duration ratio obtained with the Gaussian fit (1.414) matches the known theoretical value of $\sqrt{2}$ for a Gaussian pulse, and the experimental spectrum yields a closely matching value (1.402). The sech² fit deviates more substantially from both (1.569), consistent with its theoretical value of 1.543 for a sech pulse.

Transform-limited pulse durations in Table S3 are obtained by applying an inverse FFT to the frequency-domain spectral amplitude (with flat spectral phase) and measuring the FWHM of the resulting time-domain intensity profile; the Gaussian fit yields 34.9 fs, consistent with the analytical result from the measured spectral width ($\lambda_0 = 1284$ nm, $\Delta\lambda = 69.6$ nm, $c_B = 0.441$), while the sech² fit yields 25.2 fs. We therefore adopted the Gaussian spectral model for all simulations in this work.

**Table S3. | Simulated pulse quantities for three spectral inputs at 1300 nm.**

| Quantity | Experimental | Sech² fit | Gaussian fit |
|---|---|---|---|
| Center wavelength (nm) | ~1284 | 1284.39 | 1284.39 |
| Spectral FWHM (nm) | 69.6 | 68.7 | 69.6 |
| TL pulse duration (fs) | 32.8 | 25.2 | 34.9 |
| AC envelope FWHM (fs) | 41.6 | 33.4 | 44.6 |
| AC envelope / pulse ratio | 1.268 | 1.358 | 1.282 |
| 2nd order intensity AC / pulse ratio | 1.402 | 1.569 | 1.414 |
| NSSR (spectral fit) | -- | 0.002999 | 0.003163 |

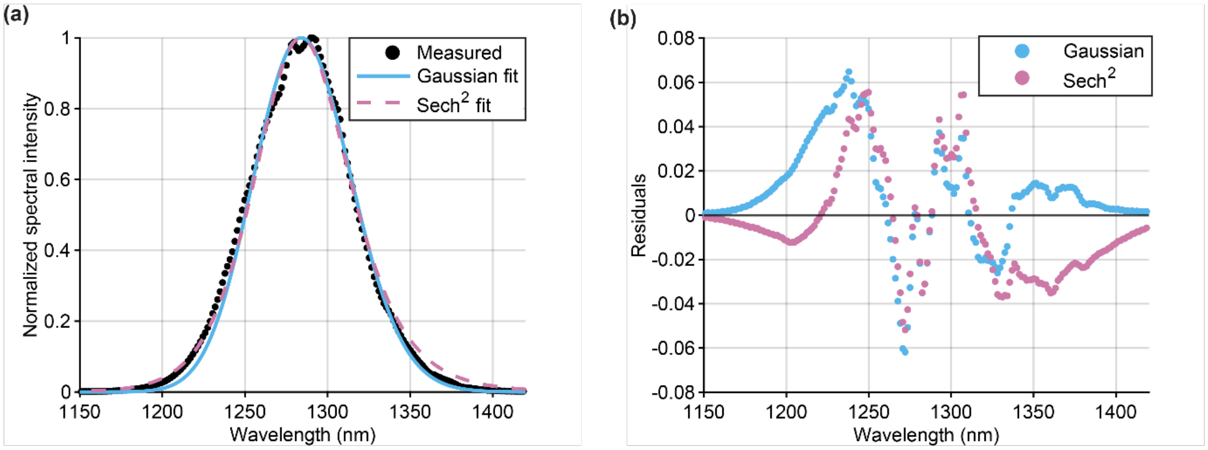


**Figure S3.** | **Spectral model selection for pulse simulations. (a)** Measured laser spectrum at 1300 nm (dots) with Gaussian (solid) and sech² (dashed) fits, both performed in the frequency domain. **(b)** Fit residuals for the Gaussian (blue) and sech² (pink) models.

A similar procedure was performed at the 1600 nm excitation window; results are presented in Table S4 and Fig. S4.

**Table S4.** | **Simulated pulse quantities for three spectral inputs at 1600 nm.**

| Quantity | Experimental | Sech² fit | Gaussian fit |
|---|---|---|---|
| Center wavelength (nm) | ~1580 | 1582.8 | 1582.8 |
| Spectral FWHM (nm) | 66.1 | 66.1 | 64.7 |
| TL pulse duration (fs) | 50.8 | 39.4 | 55.6 |
| AC envelope FWHM (fs) | 64.8 | 53.0 | 71.4 |
| AC envelope / pulse ratio | 1.276 | 1.345 | 1.284 |
| 2nd order intensity AC / pulse ratio | 1.406 | 1.533 | 1.417 |
| NSSR (spectral fit) | -- | 0.007904 | 0.007859 |

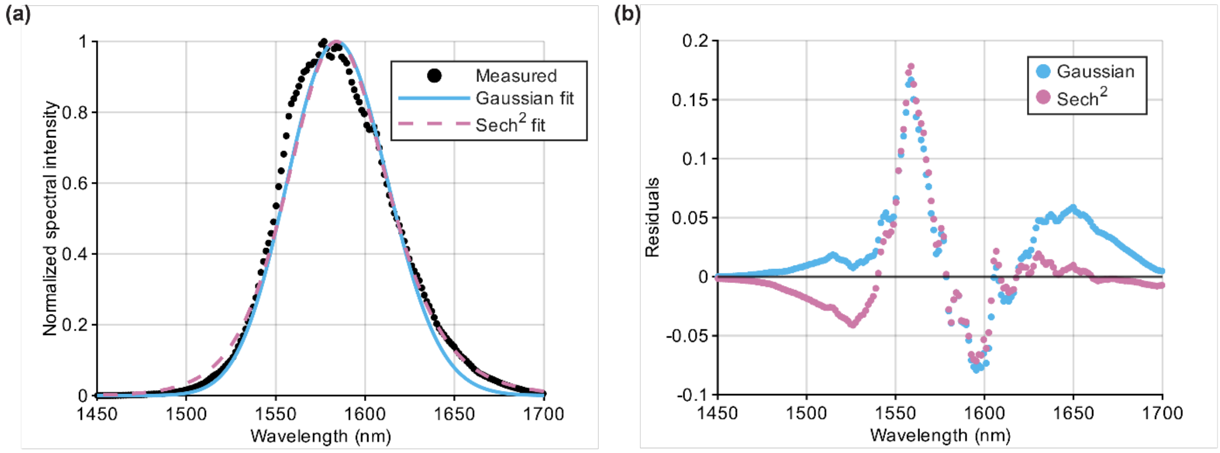


**Figure S4.** | **Spectral model selection for pulse simulations. (a)** Measured laser spectrum at 1600 nm (dots) with Gaussian (solid) and sech² (dashed) fits, both performed in the frequency domain. **(b)** Fit residuals for the Gaussian (blue) and sech² (pink) models.

### Supplementary Note S3: Data acquisition and motor control

Movement of the motorized actuator (Z812B, Thorlabs) is managed through a K-Cube Brushed DC Servo Motor Controller (KDC101, Thorlabs). A custom MATLAB class, motor.m (commit a268589)[1], wraps the Thorlabs Kinesis API (version 1.14.37)[2]. The class is used to perform basic movements with the motorized actuator needed for varying the delay time between interfering pulses.

Acquisition parameters are set before each run: motor speed, start position, and end position are manually specified in the code. These parameters must stay within the range of the motorized actuator (speed: 0.05 - 1.60 mm/s, travel range: 12 mm). The start and end positions determine the travel range of the motor, which, along with the speed, is used to calculate the travel duration. Using this duration and the repetition rate of the laser, the number of overlapping pulse pairs expected to pass through the module during this time is calculated. This number is used to configure the number of samples the vDAQ (MBF Bioscience) will acquire.

Data acquisition is managed through a separate custom class, that communicates with the vDAQ through ScanImage's connection to its Field-Programmable Gate Array (FPGA). Data acquisition is pre-configured to record a set number of samples equal to the previously calculated number of expected pulse pairs.

Once all initialization steps are complete, the motor is homed and moved to the specified start position. The motor is then set to the specified run speed and begins continuous motion, triggering the vDAQ to begin acquisition. When the number of expected samples is reached, acquisition ends and the motor is stopped.

1. Chmyrov, A. Driver for Thorlabs BBD302/PRM1Z8/K10CR1 motorized stages. (2026).
2. Kinesis Software. Thorlabs (2026).

### Supplementary Note S4: Fourth-order dispersion

Our forward model truncates the spectral-phase expansion of Eq. (4) at the third order, describing the pulse using $GDD_0$ and $TOD_0$ alone. As noted in Section 2.6.1, any fourth-order or non-polynomial spectral phase present in the actual pulse may not be accurately captured. To probe whether a fourth-order term (FOD) will improve the joint-fit (fit detailed in Supplementary Note S6), we extended the model and refit the experimental joint-measurement envelopes while holding $GDD_0$ and $TOD_0$ fixed at their recovered values. Fig. S5 shows the result for fitting $FOD_0$ to the representative 1300 nm measurement pair in Fig. 5. Adding $FOD_0$ reduces the joint-fit envelope cost by 19.6% (1.06 to 0.85), visibly improving the fit agreement with the outer wings of the envelope. Notably, the 1-rod measurement widened to a pulse FWHM of 72 fs (previously 58 fs with $FOD_0$ = 0 $fs^4$). The recovered $FOD_0$ was consistent across the nine measurement permutations, with a relative standard deviation of 20% (Laser 1) and 14% (Laser 2) and differed between the two lasers: $FOD_0$ = 850,000 ± 170,000 $fs^4$ [mean ± std] (Laser 1) and 243,000 ± 35,000 $fs^4$ (Laser 2), at the GDD-minimized compressor setting.

While adding an FOD term improves our envelope fit, we cannot determine the underlying cause of the improvement. It is consistent with a fourth-order spectral phase in a pulse, but an additional free parameter can equally absorb residual trace asymmetry, measurement noise, an incorrectly identified carrier frequency, or non-polynomial phase that our GDD/TOD forward model does not capture.

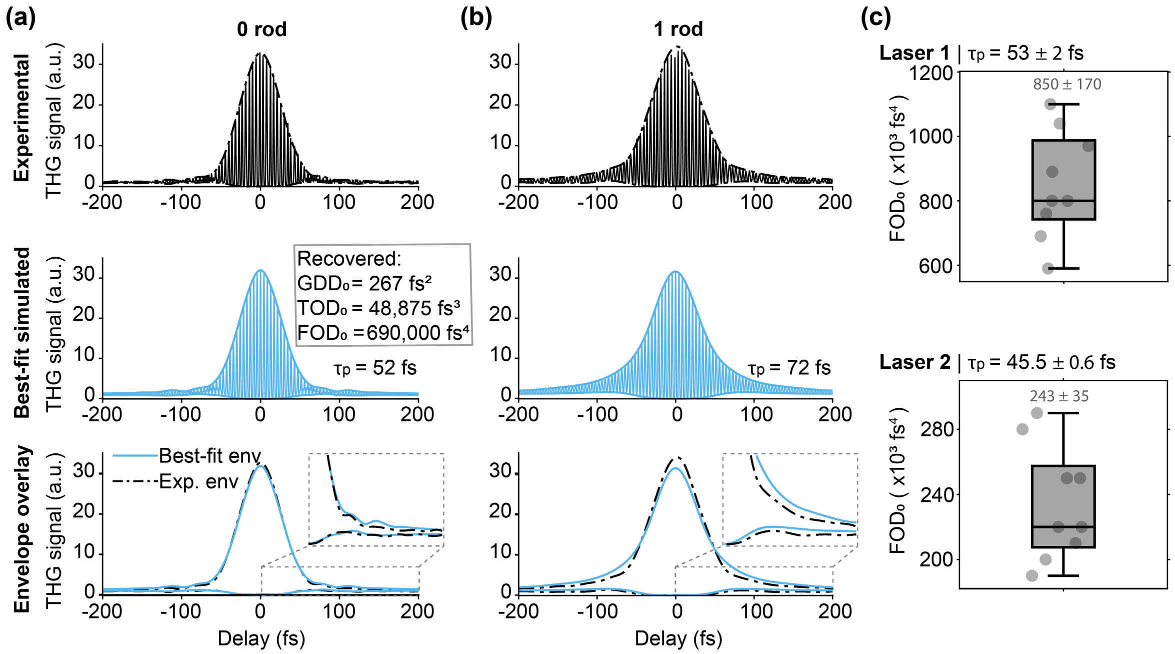


**Figure S5. | Dispersion estimation from 1300 nm THG interferometric autocorrelations after fitting $FOD_0$ with fixed $GDD_0$ and $TOD_0$.** Starting from the joint-fit of Fig. 5, $GDD_0$ and $TOD_0$ were held fixed at their recovered values and a fourth-order dispersion term ($FOD_0$) was added as the only free parameter. **(a,b)** Representative 0-rod **(a)** and 1-rod **(b)** experimental THG-IAC traces at the optimal compressor setting on Laser 1, with best-fit simulated traces and envelope overlays. For this representative pair, the recovered fourth-order dispersion was $FOD_0$ = 690,000 $fs^4$ (with $GDD_0$ = 267 $fs^2$, $TOD_0$ = 48,875 $fs^3$). This corresponds to a pulse duration of $\tau_P$ = 52 fs (0-rod) and 72 fs (1-rod). **(c)** Across N = 9 permutations of the measured traces, the recovered fourth-order dispersion was $FOD_0$ = 850,000 ± 170,000 $fs^4$ [mean ± std] for Laser 1 and 243,000 ± 35,000 $fs^4$ for Laser 2 (same laser model), at their respective optimal settings. The corresponding baseline pulse durations for the permutations were $\tau_P$ = 53 ± 2 fs (Laser 1) and 45.5 ± 0.6 fs (Laser 2).

### Supplementary Note S5: Dispersion Look-Up-Table Estimation (D-LUTE)

Dispersion Look-Up-Table Estimation (D-LUTE) is an initialization method for the joint-measurement THG-IAC, $G^{(3)}(\tau)$, fitting procedure used to estimate baseline pulse dispersion. Because autocorrelation-based recovery can be susceptible to local minima without good initial starting points, D-LUTE first identifies plausible baseline-dispersion regions using a simple pulse observable that can be measured experimentally and numerically predicted.

In this work, the D-LUTE observable is the FWHM of the intensity-autocorrelation component, $A^{(3)}(\tau)$, extracted from each THG-IAC trace. The width is used because it is a single, readily obtainable value from the same processed traces used in the full fitting pipeline. The same framework could be extended to other observables, such as 2P or 3P fluorescence signal metrics, provided that the observable's response to added dispersion differs across the relevant GDD/TOD parameter space.

D-LUTE uses a precomputed lookup table, $F_{LUT}(\varphi_2, \varphi_3)$, that relates total GDD/TOD values to the expected intensity autocorrelation FWHM. The unknown baseline dispersion is denoted $(\varphi_{2,0}, \varphi_{3,0})$, corresponding to $GDD_0$ and $TOD_0$. For measurement $m$, the inserted dispersive material contributes known dispersion offsets. After accounting for these offsets, D-LUTE evaluates candidate baseline values by asking which $(\varphi_{2,0}, \varphi_{3,0})$ best explain the measured FWHMs, $\gamma_m$. The following sections describe how the LUT is generated, how measurement-specific likelihood surfaces are calculated and combined, and how candidate initialization points are selected.

#### Look-up table generation

To implement D-LUTE on our 1300 nm pulses, a numerical look-up table (LUT) was first generated to relate dispersion values, $(\varphi_2, \varphi_3)$, to the FWHM of the corresponding intensity-autocorrelation trace. The LUT was computed using the estimated transform-limited pulse duration $\tau_{P,TL}$, estimated from the measured pulse spectrum.

For a given pulse spectrum $\tilde{E}(\omega)$, the LUT was generated by numerically simulating $G^{(3)}(\tau)$ traces over a range of dispersion values. Specifically, $\varphi_2$ (GDD) was varied from $-$ 9 kfs$^2$ to + 9 kfs$^2$ range in 100 fs$^2$ steps, and $\varphi_3$ (TOD) from $-$ 160 kfs$^3$ to + 220 kfs$^3$ in 2 kfs$^3$ steps. Each simulation covered a 4 ps delay window with 0.7 fs temporal resolution. The zeroth- and first-order dispersion terms, $\varphi_0$ and $\varphi_1$, were omitted since they only introduce a constant phase shift and a temporal offset, respectively, and do not affect pulse shape.

For each $(\varphi_2, \varphi_3)$ pair, the dispersed time-domain electric field $E(t)$ was obtained via inverse Fourier transform and used to simulate the third-order autocorrelation trace, $G^{(3)}(\tau)$. The simulated trace was then processed in the same way as the experimental data. $G^{(3)}(\tau)$ was spectrally filtered to isolate the low-frequency autocorrelation component, $A^{(3)}(\tau)$. Although the intensity autocorrelation can be calculated directly from $E(t)$, filtering the simulated THG-IAC reflected the experimental procedure used to extract the observable. The filtered trace was then normalized, bias-corrected by subtracting one, and its FWHM, $\gamma$, recorded in the LUT alongside the corresponding dispersion values $\varphi_2$ and $\varphi_3$ as $F_{LUT}(\varphi_2, \varphi_3)$.

#### Likelihood mapping

To estimate the base system dispersion, denoted $GDD_0 = \varphi_{2,0}$ and $TOD_0 = \varphi_{3,0}$, D-LUTE constructed a likelihood surface over dispersion values $(\varphi_2, \varphi_3)$ by comparing measured and simulated intensity autocorrelation FWHM values. For each measured autocorrelation trace, corresponding to a known amount of added GDD and TOD $(\varrho_m, \zeta_m)$, an offset LUT surface $(F_m)$ was generated from the baseline LUT, $F_{LUT}$, as:

$$F_m(\varphi_2, \varphi_3) = F_{LUT}(\varphi_2 - \varrho_m, \varphi_3 - \zeta_m).$$

A Gaussian likelihood surface, $P_m$, was then calculated for each measured FWHM $\gamma_m$ using:

$$P_m(\varphi_2, \varphi_3) = e^{-\frac{(F_m(\varphi_2,\varphi_3)-\gamma_m)^2}{2\sigma_{\gamma_m}^2}}.$$

The uncertainty $\sigma_{\gamma_m}$ was set to 15% of $\gamma_m$. The overall likelihood map was then obtained by multiplying individual surfaces across all measurements:

$$P_{total}(\varphi_2, \varphi_3) = \prod_m P_m(\varphi_2, \varphi_3).$$

While only showing likelihood values above a threshold of 0.3, Fig. S6 shows how two likelihood surfaces combined to produce $P_{total}$, with a constrained region of high likelihood. This surface is plotted on top of the utilized LUT.

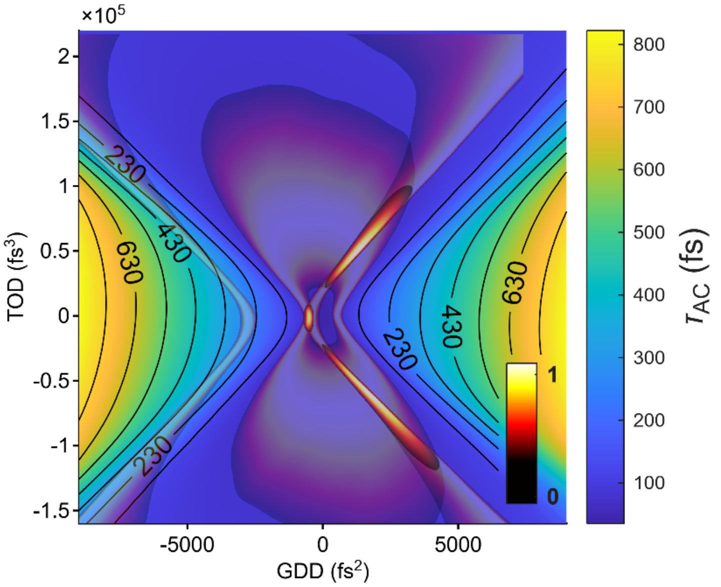


**Figure S6. | D-LUTE likelihood surfaces for a set of measurements shown over the utilized LUT.** The underlying contour map shows the simulated intensity-autocorrelation FWHM from the precomputed LUT. Superimposed on this, semi-transparent colored bands indicate regions consistent with the measured FWHM values for individual rod conditions after accounting for their known added dispersion. The overlap of these shifted likelihood surfaces produces the combined high-likelihood region used to identify candidate baseline dispersion values for initialization of the joint THG-IAC fit. The likelihood value is indicated by the inset colorbar.

After the total likelihood surface, $P_{total}(\varphi_2, \varphi_3)$, was calculated, D-LUTE identifies candidate baseline-dispersion values from regions of high likelihood. The likelihood surface was first normalized and thresholded (> 0.3) to retain only candidate regions that strongly supported the measured FWHM values. These high-likelihood regions were then clustered, and the peak-likelihood point within each cluster was selected as a candidate ($GDD_0$, $TOD_0$) initialization point. Each selected D-LUTE candidate was subsequently evaluated by first spectrally filtering the corresponding autocorrelations to isolate their $2\omega_0$ components, then computing the envelope cost of Eq. (9) on these filtered traces. The candidate with the lowest cost was carried forward as the starting point for the joint-fit procedure. We found this $2\omega_0$ spectral component isolation necessary for robust candidate selection. The cost function evaluated on the unfiltered envelopes occasionally converged to the origin when significant noise was present in the measurements. In contrast, the $2\omega_0$ spectral component, which we observed to be sensitive to TOD, yielded more noise-robust selection.

### Limitations

D-LUTE assumes that the lookup-table spectrum and GDD/TOD pulse model are appropriate for the measured pulse, and requires that the selected rod conditions contribute a significant amount of dispersion to a pulse. The N-SF11 rods contribute significant GDD, allowing D-LUTE to promote the correctly signed GDD value. However, the small amount of added TOD hinders the procedure's ability to identify the correctly signed $TOD_0$. Adding more TOD would further reduce the likelihood of the incorrect ambiguous pulse dispersion. As mentioned in Section 2.6.4, we circumvented the $TOD_0$ sign ambiguity by assuming $TOD_0$ is positive.

### Supplementary Note S6: Joint-measurement fit procedure implementation

The two-staged fit procedure introduced in Section 2.6.3 recovers the baseline second-order and third-order dispersion ($GDD_0$, $TOD_0$) by jointly fitting the 0- and 1-rod THG-IAC trace envelopes to the forward model (detailed in Section 2.6.1). For each rod condition, the simulated trace is evaluated at the total dispersion ($GDD_0 + m\ GDD_{N\text{-}SF11}$, $TOD_0 + m\ TOD_{N\text{-}SF11}$), with the N-SF11 rod increments held fixed, and scored against experiment by the envelope cost $L$ of Eq. (9). The joint cost is the sum over the two rod conditions, and the fit minimizes it in two stages initialized at the best of the D-LUTE candidate estimate. Before fitting, the zero-delay position of each measured trace is aligned by fitting a Gaussian to its intensity autocorrelation and taking the fitted center as delay = 0 fs.

*Stage 1 – grid search.* A constrained grid search first localizes the cost minimum on a grid centered on the D-LUTE estimate ($GDD_0^{init}$, $TOD_0^{init}$), with fixed steps in each axis:

$$GDD_0 \in [GDD_0^{init} - \Delta G,\ GDD_0^{init} + \Delta G], \quad TOD_0 \in [TOD_0^{init} - \Delta T,\ TOD_0^{init} + \Delta T\ ]$$

At 1300 nm we jointly fit the 0- and 1-rod traces with $\Delta G = 500$ fs$^2$, $\Delta T = 20{,}000$ fs$^3$ (a 41 × 41 node grid). At 1600 nm we fit the 0- and 3-rod traces with $\Delta G = 400$ fs$^2$, $\Delta T = 30{,}000$ fs$^3$ (a 33 × 61 node grid). Both used steps of 25 fs$^2$ in GDD and 1000 fs$^3$ in TOD, and envelopes are scored over $|\tau| \leq W = 1100$ fs. The joint envelope cost is evaluated at every node, and the lowest-cost node ($GDD_0^*$, $TOD_0^*$) is passed to Stage 2. Due to the window sizes, a good starting estimate is important. Poor initializations are corrected by the boundary check described below.

*Stage 2 – least-squares refinement.* Starting from the grid optimum identified in Stage 1, the parameters are refined by nonlinear least squares (MATLAB *lsqnonlin*) minimization of the joint-fit cost. Since $GDD_0$ and $TOD_0$ differ by orders of magnitude, the solver optimizes dimensionless multipliers ($m_{GDD}$, $m_{TOD}$) about the grid optimum,

$$(GDD_0, TOD_0) = (GDD_0^*, TOD_0^*) \bullet (m_{GDD}, m_{TOD})$$

As the original Stage 1 grid window is narrow and centered on the D-LUTE estimate, a poor initialization can place the true minimum outside it. We perform a grid check on the refined value from the nonlinear least squares search, by checking if the value is near (within two steps) or beyond the grid search boundaries. If this is the case, the grid is recentered on the refined dispersion values and stages 1-2 are repeated. This loop iterates until the refined optimum lies within the interior, walking the search toward the true minimum in the event of a poor initialization.

### Supplementary Note S7: Cost-function preference for positive $TOD_0$ in the joint-measurement fit.

Jointly fitting the 0-rod and 1-rod traces reduces the degeneracy described in Section 2.6.4, as inserting the N-SF11 rod affects the magnitude of the negative and positive dispersion terms differently (Fig. 3c). As the N-SF11 rods add little TOD relative to GDD at 1300 nm, the wrong-signed TOD candidate is suppressed but not eliminated. For the example measurement trace shown in Fig. 5a-b, we removed the TOD > 0 constraint and had D-LUTE return candidates in both positive and negative TOD regions. These candidates were then refined independently through the full joint-fit (Table S5). The positive-$TOD_0$ candidate produced a lower joint-measurement cost during D-LUTE candidate selection (0.36 vs 0.41), as well as after the joint-fit (1.06 vs 1.42). The N-SF11 rods add positive TOD, and the cost is lowered when the $TOD_0$ magnitude is increased between the 0-rod and 1-rod measurements. Thus, the positive $TOD_0$ is promoted over negative $TOD_0$. Overall, the joint-fit cost favored the positive $TOD_0$ estimate across all GDD-optimized measurement pair permutations tested (9/9 for Laser 1, 9/9 for Laser 2).

Together with the positive residual TOD reported for similar laser sources and the positive TOD expected from the microscope's material dispersion at 1300 nm, it motivates the $TOD_0 > 0$ constraint (Section 2.6.4). The constraint concerns only the recovered sign. As shown in Fig. S9 (Supplementary Note S9), the modeled pulse duration and THG signal are approximately symmetric with respect to the sign of $TOD_0$. A negative $TOD_0$ of equal magnitude would produce comparable pulse broadening and signal loss, differing only in the sign of the required compensation.

**Table S5. | Joint-fit cost comparison for positive- and negative- $TOD_0$ candidates for Laser 1, during both candidate selection and final joint-fit.**

| Branch | Candidate $GDD_0$ | Candidate $TOD_0$ | Candidate 2ω cost | Final $GDD_0$ | Final $TOD_0$ | Final ENV cost |
|---|---|---|---|---|---|---|
| Positive TOD | 910 $fs^2$ | 53200 $fs^3$ | 0.36 | 267 $fs^2$ | 48900 $fs^3$ | 1.06 |
| Negative TOD | 1220 $fs^2$ | -58400 $fs^3$ | 0.41 | 151 $fs^2$ | -32600 $fs^3$ | 1.42 |

### Supplementary Note S8: Estimated dispersion at different settings of the Cronus 3P laser's internal prism compressor

The Cronus 3P laser contains an internal prism compressor that can be used to compensate for GDD and minimize the pulse duration at the microscope focal plane. To validate the GDD estimates and identify the compressor setting that produced the lowest residual dispersion for imaging, we acquired measurement pairs across a range of compressor settings at both 1300 nm and 1600 nm windows.

At the 1300 nm window, three measurements were acquired without glass rods and three measurements were acquired with one glass rod inserted at each compressor setting. GDD and TOD were then estimated by fitting all nine permutated combinations of these measurements. Compressor compensation was varied from -4.6 to -5.8 $kfs^2$ in 0.4 $kfs^2$ increments, with an additional measurement acquired at the laser's maximum negative compensation setting of -6.0 $kfs^2$. The resulting dispersion estimates are reported in Table S6 and plotted in Fig. S7.

As shown in Fig. S7a and Table S6, increasing the magnitude of negative compressor compensation reduced the estimated residual GDD at the focal plane, with the GDD crossing zero between -5.4 and -5.8 $kfs^2$. In contrast, TOD remained positive across all compressor settings and varied less systematically with compressor compensation (Fig. S7b, Table S6), indicating that the compressor cannot be efficiently utilized for joint compensation of both GDD and TOD.

**Table S6. | Estimated 1300 nm window residual dispersion at the microscope focal plane for different prism compressor settings.**

| Compressor Compensation ($kfs^2$) | Estimated GDD ($fs^2$) | Estimated TOD ($fs^3$) |
|---|---|---|
| -6.0 | -441 ± 261 | 60200 ± 5800 |
| -5.8 | -75 ± 206 | 65600 ± 3000 |
| -5.4 | 289 ± 99 | 54400 ± 4900 |
| -5.0 | 549 ± 63 | 65100 ± 3000 |
| -4.6 | 1019 ± 129 | 46900 ± 12100 |

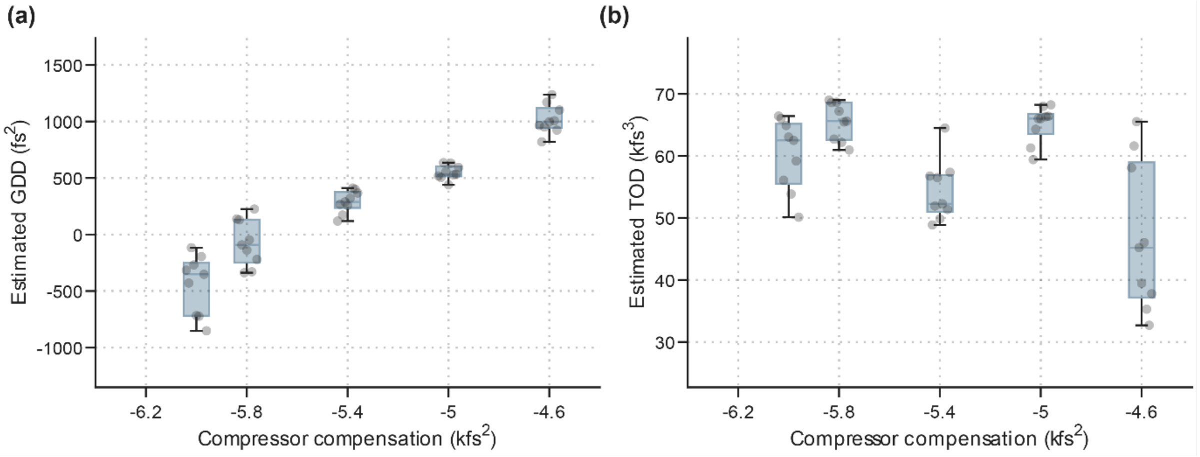


**Figure S7. | Estimated 1300 nm pulse dispersion with different prism compressor compensation.** Estimated **(a)** GDD and **(b)** TOD of 1300 nm pulses with different amounts of GDD compensation applied by the laser's internal prism compressor. Gray points show estimated values from different measurement pairs, with blue box plots summarizing the distribution for each compressor setting. Points are horizontally jittered to improve visibility.

We repeated the prism compressor optimization near 1600 nm to determine the compressor setting that minimized residual dispersion at the microscope focal plane. Compressor compensation was varied from 2.0 to 7.0 $kfs^2$ in 0.5 $kfs^2$ increments. The resulting estimates showed that residual GDD increased from negative to positive values across this range, crossing zero between 5 and 5.5 kfs². Residual TOD across all compressor settings was higher than at the 1300 nm window, but exhibited a similar insensitivity to compressor GDD compensation. These 1600 nm dispersion estimates are reported in Table S7 and plotted in Fig. S8.

**Table S7. | Estimated 1600 nm window residual dispersion at the microscope focal plane for different prism compressor settings.**

| Compressor Compensation (kfs²) | Estimated GDD (fs²) | Estimated TOD (fs³) |
|---|---|---|
| 2.0 | -3350 ± 62 | 105100 ± 3400 |
| 2.5 | -3200 ± 206 | 117600 ± 19900 |
| 3.0 | -2367 ± 229 | 141200 ± 33900 |
| 3.5 | -2282 ± 47 | 104800 ± 7100 |
| 4.0 | -1001 ± 462 | 130300 ± 4200 |
| 4.5 | -1141 ± 322 | 118200 ± 3500 |
| 5.0 | -273 ± 558 | 121400 ± 12900 |
| 5.5 | 669 ± 278 | 138900 ± 7800 |
| 6.0 | 1403 ± 293 | 117900 ± 40200 |
| 6.5 | 1670 ± 175 | 78100 ± 28800 |
| 7.0 | 1716 ± 233 | 64600 ± 42000 |

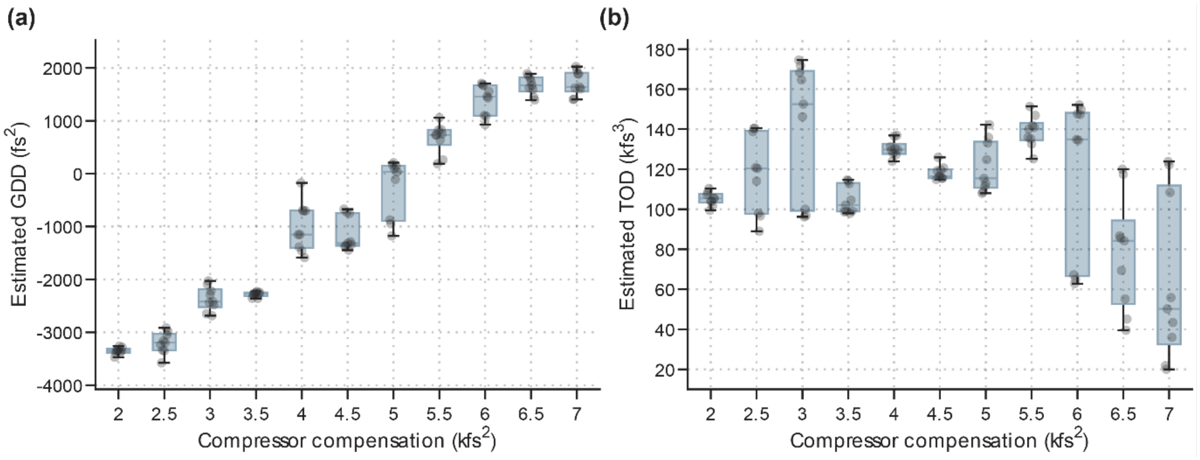


**Figure S8. | Estimated 1600 nm pulse dispersion with different prism compressor compensation.** Estimated **(a)** GDD and **(b)** TOD of 1600 nm pulses with different amounts of GDD compensation applied by the laser's internal prism compressor. Gray points show estimated values from different measurement pairs, with blue box plots summarizing the distribution for each compressor setting. Points are horizontally jittered to improve visibility.

The values in Tables S6 and S7 are those measured with the autocorrelator module in the beam path. Subtracting the BK7 beamsplitter's double-pass contribution (Section 2.2) gives the residual GDD present during standard imaging, which is smallest at −5.4 kfs² at 1300 nm and at 4.5 kfs² at 1600 nm – the settings used in Sections 3.2 and 3.3.

### Supplementary Note S9: Effect of pulse dispersion on third-harmonic generation (THG) signal

Because third-harmonic generation (THG) scales with the cube of instantaneous intensity, the total THG signal generated at the focal point depends on the pulse's temporal profile rather than its energy alone. For a pulse with fixed energy, the generated THG signal is proportional to $\int I^3(t;\varphi_2,\varphi_3)dt$, where $I(t;\varphi_2,\varphi_3) = |E(t;\varphi_2,\varphi_3)|^2$ is provided by Eq. (5). We define the signal fraction, $\eta(\varphi_2,\varphi_3)$, as the THG signal generated by a dispersed pulse normalized to that of the transform-limited pulse:

$$\eta(\varphi_2,\varphi_3) = \frac{\int I^3(t;\varphi_2,\varphi_3)dt}{\int I^3(t;0,0)dt} \quad \text{(S1)}$$

If dispersion stretches the pulse without altering its envelope shape, the peak intensity falls as $1/\tau_p$ at fixed pulse energy, so the time-integrated THG signal, $\int I^3(t)dt$, scales as $(1/\tau_p)^3 \cdot \tau_p = 1/\tau_p{}^2$, and Eq. (S1) reduces to Eq. (S2). Here, $\tau_{p,TL}$ is the pulse duration of the transform-limited pulse.

$$\eta = \left(\frac{\tau_{p,TL}}{\tau_p}\right)^2 \quad \text{(S2)}$$

Equation (S2) is exact for Gaussian pulses with only GDD, since GDD broadens a Gaussian pulse while preserving its envelope shape. TOD, however, alters the temporal envelope by redistributing energy from the central pulse into temporal side lobes. Consequently, the reduction in signal is greater than would be predicted from pulse broadening alone, and the FWHM of the central pulse no longer uniquely determines the THG signal. As a result, Eq. (S2) systematically overestimates the signal fraction for pulses with significant TOD.

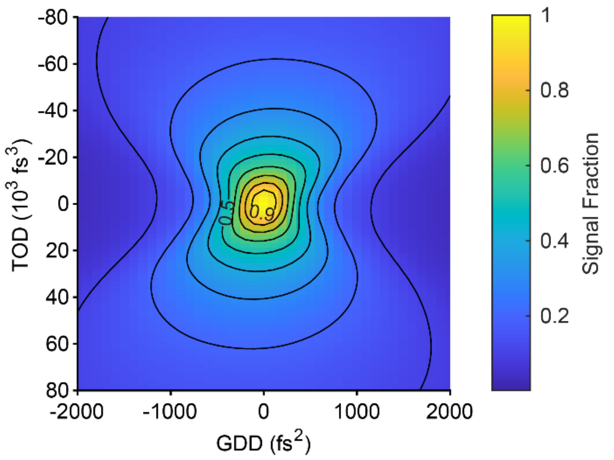


**Figure S9. | Simulated third-harmonic signal fraction as a function of residual dispersion.** Signal fraction η (Eq. S1) for a simulated 1300 nm pulse (Laser 1, $\lambda_0$ = 1284 nm, $\Delta\lambda$ = 69.6 nm , $\tau_{p,TL}$ = 32.8 fs) over a grid of GDD and TOD, normalized to the transform-limited pulse.

Fig. S9 maps the calculated THG signal fraction for a simulated pulse from our Laser 1 1300 nm spectrum ($\lambda_0 = 1284\ nm$, $\Delta\lambda = 69.6\ nm$ , $\tau_{p,TL} = 32.8\ fs$), over a range of GDD and TOD values. At the selected prism-compressor setting, the recovered baseline dispersion with the autocorrelator module inserted was $GDD_0$ = 289 fs$^2$ and $TOD_0$ = 54,400 fs$^3$. A pulse with this residual dispersion has a pulse duration of $\tau_p = 55$ fs and generates a signal fraction $\eta = 0.219$. The common method of estimating the effect of dispersion using pulse width alone, Eq. (S2), expects $\eta_{S2} = 0.36$. This deviation is due to how TOD not only widens the pulse, but also redistributes energy away from the central pulse lobe. Setting $GDD_0$ to zero while retaining $TOD_0$ = 54,400 fs$^3$ marginally raises the signal fraction to 0.227, demonstrating that residual TOD is the dominant limitation.

The corresponding dispersion during standard imaging, with the autocorrelator module bypassed, is estimated by subtracting the BK7 beamsplitter's calculated double-pass contribution from the fitted baseline values. At 1284 nm, subtracting 139 $fs^2$ of GDD and 3237 $fs^3$ of TOD gives $GDD_0$ = 150 $fs^2$ and $TOD_0$ = 51,163 $fs^3$. Evaluating Eq. (S1) at these values gives $\eta = 0.237$. Within the GDD/TOD model, full compensation would therefore increase the signal by ~4.2× at fixed pulse energy, or equivalently reduce the required power by 38% for equal signal levels.
